# Dynamic mitochondrial transcription and translation in B cells control germinal centre entry and lymphomagenesis

**DOI:** 10.1101/2022.07.19.500689

**Authors:** Yavuz F Yazicioglu, Eros M Marin, Ciaran Sandhu, Silvia Galiani, Iwan G A Raza, Mohammad Ali, Barbara Kronsteiner, Ewoud B Compeer, Moustafa Attar, Susanna J Dunachie, Michael L Dustin, Alexander J Clarke

## Abstract

Germinal centre (GC) B cells undergo proliferation at very high rates in a hypoxic microenvironment, but the cellular processes driving this are incompletely understood. Here we show that the mitochondria of GC B cells are highly dynamic, with significantly upregulated transcription and translation rates associated with the activity of transcription factor mitochondrial A (TFAM). TFAM, whilst also necessary for normal B cell development, is required for entry of activated GC-precursor B cells into the germinal centre reaction, and deletion of *Tfam* significantly impairs GC formation, function, and output. Loss of TFAM in B cells compromises the actin cytoskeleton and impairs cellular motility of GC B cells in response to chemokine signalling, leading to their spatial disorganisation. We show that B cell lymphoma substantially increases mitochondrial translation, and deletion of *Tfam* in B cells is protective against the development of lymphoma in a c-Myc transgenic model. Finally, we show that pharmacologic inhibition of mitochondrial transcription and translation inhibits growth of GC-derived human lymphoma cells, and induces similar defects in the actin cytoskeleton.

## Introduction

The germinal centre (GC) reaction is a highly spatially organised process in secondary lymphoid tissue essential for humoral immunity^1^. B cells responding to antigen captured by follicular dendritic cells introduce random mutations in their immunoglobulin genes in a process known as somatic hypermutation (SHM), which occurs in the anatomically defined dark zone (DZ), and are then selected through competitive interaction with follicular T-helper cells (T_FH_) in the light zone (LZ). GC B cells cycle between these zones, leading to antibody affinity maturation and eventually formation of memory B or plasma cells. During SHM, mutations are introduced in immunoglobulin gene loci through the action of activation-induced cytosine deaminase (AICDA). This can give rise to oncogenic mutations, for example the translocation of MYC with *IGH* or *IGL* loci^2^. GC B cells are the origin of most diffuse large B cell lymphomas (DLBCL), the most common non-Hodgkin lymphoma.

The metabolic processes that support GC B cell homeostasis remain incompletely understood. GC B cells are highly proliferative, with division times as short as 4-6 hours, and reside within a hypoxic microenvironment^3–5^. Typically, rapidly proliferating immune cells mainly use aerobic glycolysis, but GC B cells rely on fatty acid oxidation and oxidative phosphorylation (OxPhos), although to what extent remains somewhat controversial^4–10^. This metabolic phenotype is carried over into a substantial proportion of diffuse large B cell lymphoma (DLBCL), which are also OxPhos dependent^11^. Mitochondria have been demonstrated in vitro to be regulators of B cell signalling through redox-related mechanisms, although whether these studies can be generalised to the hypoxic microenvironment of the GC found in vivo is uncertain^12–14^.

Regulation of mitochondrial transcription and translation has been shown to be important in other immune cell types, including cytotoxic CD8^+^ T cells, where it controls cell killing independent of effects on metabolism, and cytokine production in CD4^+^ T cells^15–17^. A key regulator of mitochondrial transcription and translation is *transcription factor A mitochondrial* (TFAM), a DNA-binding high mobility box group (HMBG) protein which aids in the packaging of the mitochondrial genome into nucleoids, analogous to the role of histone proteins, and also controls transcription and translation of mitochondrial DNA, serving as a regulator of mitochondrial biogenesis^18,19^. In T cells, TFAM controls immunosenescence by restraining the production of inflammatory cytokines^20^, and in fibroblasts and myeloid cells acts to regulate an anti-viral immune state^21^. Recent work has shown that the mitochondrial DNA helicase TWINKLE is required for GC formation and plasma cell differentiation, but is dispensable for B cell development^22^. However, the dynamics, function, and regulation of mitochondria in GC B cell biology remains incompletely understood.

We show that GC B cell mitochondria are highly dynamic organelles, undergoing profound structural changes as they transition through the germinal centre reaction. We find that TFAM is dynamically regulated in B cells, and is required for their development, and transcriptional and spatial entry into the GC reaction by modulating cellular motility. We also demonstrate using genetically modified mouse models that TFAM is essential for the development of lymphoma, and that pharmacologic inhibition of mitochondrial transcription and translation in human lymphoma cells represents a potential treatment target for human disease.

## Results

### 1. GC B cells undergo extensive mitochondrial remodelling and biogenesis associated with mitochondrial protein transcription and translation

In order to determine mitochondrial density and structure in GC B cells, we first immunised mitochondrial reporter mice expressing GFP and mCherry tagged with a mitochondrial fission-1 (FIS1) targeting sequence (MitoQC)^23^, with the T-dependent antigen sheep red blood cells (SRBC), and examined splenic tissue by confocal microscopy and flow cytometry at day 12. GFP signal was strongly localised to GL-7^+^ GCs, and significantly higher than the surrounding B cell follicle (Fig. 1A-B), which we confirmed by flow cytometry (Fig. 1C). GC B cells cycle between anatomically defined regions known as the light zone (LZ), in which T cell interaction occurs, and the dark zone (DZ), the site of rapid proliferation and somatic hypermutation. The recently identified grey zone (GZ) represents an intermediate state associated with the G2-M cell cycle phase and expression of metabolism and DNA replication-related genes^24^. Using flow cytometry, we found that the mitochondrial GFP signal was highest in the GZ, followed by the LZ and then the DZ (Fig. 1C). To examine GC B cell mitochondrial morphology in these regions, we sorted naïve B cells from unimmunised, and LZ, DZ, and GZ GC B cell subsets respectively from MitoQC mice immunised with SRBC (day 12) (for gating strategy see Extended Data Fig. 1A), and performed 3D Airyscan confocal imaging. We found highly distinct mitochondrial morphology between naive and GC B cells, with small, fragmented mitochondria in naïve B cells, and large fused mitochondria in GC B cells, most prominently in those from the GZ (Fig. 1D). Mitochondrial mass assessed by 3D volume analysis was also highest in GZ B cells, followed by LZ and then DZ cells, corresponding with flow cytometry measurements. The MitoQC reporter mouse also allows detection of autophagy of mitochondria (mitophagy), as mCherry is resistant to quenching in the acid environment of the autolysosome. Using multispectral imaging flow cytometry, we screened GC B cells for the presence of GFP^−^ mCherry^+^ punctae. Although autophagy has been reported in GC B cells^25^, we did not identify evidence of active mitophagy (Extended Data Fig. 1B-D). Thirteen protein members of the electron transport chain (ETC) are encoded by mitochondrial DNA and synthesised by the mitochondrial translation machinery. We hypothesised that in order for B cells entering the GC reaction to acquire the high mitochondrial mass we observed, activation of mitochondrial transcription and translation would be required. Using Aicda-Cre × Rosa26^STOP^tdTomato reporter mice to label GC B cells, we found markedly increased levels of cytochrome c oxidase subunit 1 (COX I), part of ETC complex IV, in the GC (Fig. 1E). This was confirmed by intracellular flow cytometry, which revealed a substantial increase in COX I levels in GC B cells, compared to IgD^+^GL-7^−^ naïve B cells (Fig. 1F).

**Figure 1.**
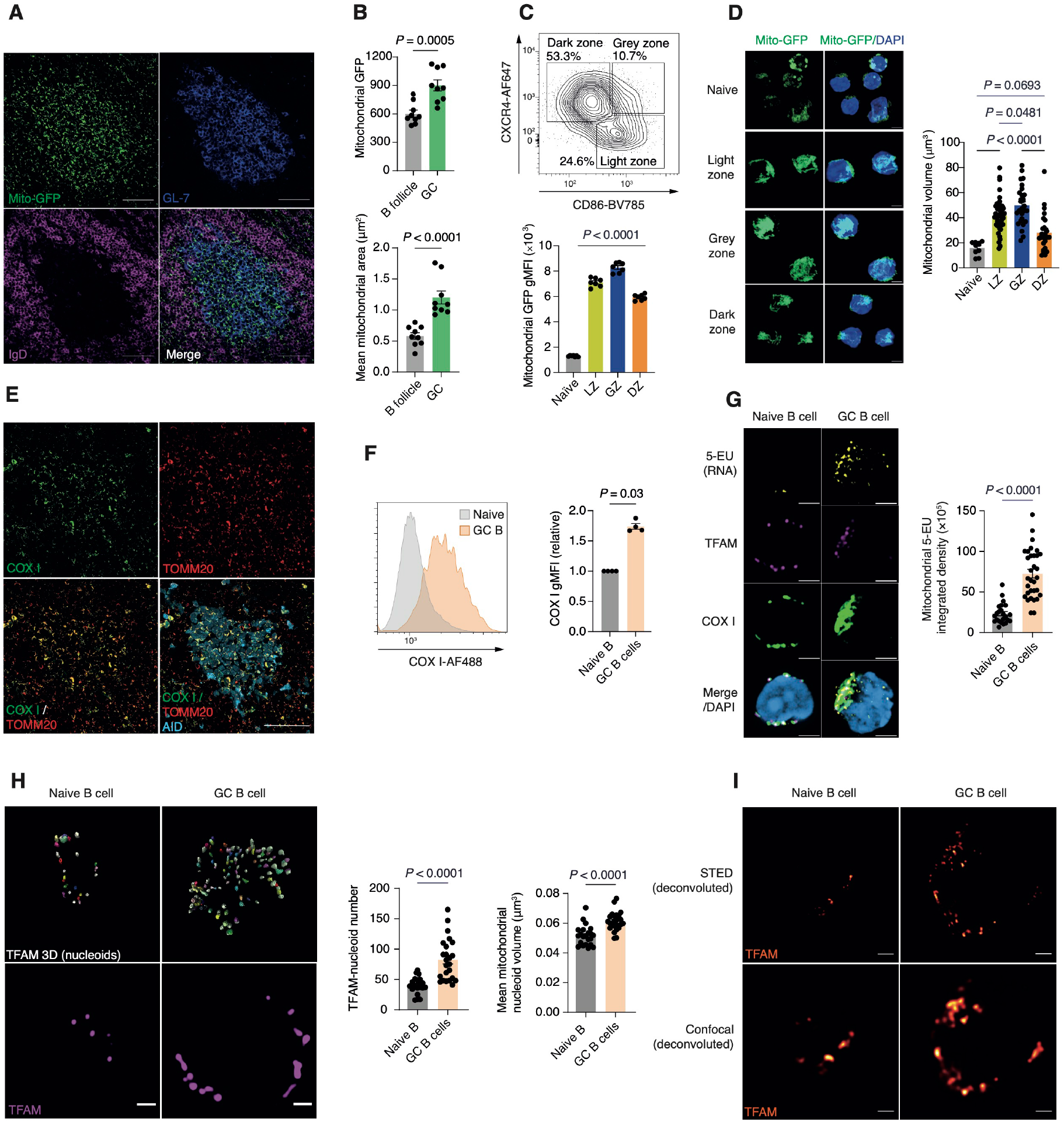
GC B cells undergo extensive mitochondrial remodelling and biogenesis associated with mitochondrial protein translation. A. Spleen sections from MitoQC mice immunised with SRBC and analysed at D12. Scale bar 50μm. Representative of three independent experiments B. Quantification of mitochondrial GFP signal intensity and area (μm^2^) in GCs. Each point represents an individual GC and surrounding B cell follicle, pooled from three individual MitoQC mice immunised as in A. Representative of two independent experiments. C. Gating strategy for GC B cells and their subsets in the spleen. GC and naïve B cells were gated as CD19^+^CD38^−^GL-7^+^ and CD19^+^IgD^+^GL-7^−^ respectively. GC B cells from the dark zone (DZ), grey zone (GZ), and light zone (LZ) were identified based on CXCR4 and CD86 expression signatures as depicted. Quantification and comparison of gMFI of mitochondrial GFP in splenic naïve, LZ, GZ and DZ GC B cells from immunised MitoQC mice (n=7), pooled from two independent experiments. D. 3D Airyscan immunocytochemistry (ICC) images depicting the mitochondrial network (GFP) in MACS-enriched and flow sorted GC B cells (LZ, GZ, and DZ) and magnetic bead sorted naïve B cells harvested from immunised (enhanced SRBC immunisation protocol at D12) and unimmunised MitoQC spleens, scale bar 3μm. 3D mitochondrial volume (μm^3^) quantification in naïve, LZ, GZ, and DZ compartments, isolated as in D. Each symbol represents one cell pooled from n=4. Representative of two independent experiments. E. Spleen sections from Aicda-Cre × Rosa26^STOP^tdTomato mice immunised with SRBC and analysed at D12, with IHC for COX I and TOMM20. Scale bar 50μm. Representative of two independent experiments. F. Flow cytometry histogram plots and quantification of COX I protein levels in GC B cells normalised to IgD^+^ naïve B cells from the same mice (n=4). Representative of three independent experiments. G. Airyscan images of in vivo 5-EU incorporation (indicating RNA synthesis), with COX I and TFAM ICC in naïve and GC B cells. Quantification of mitochondrial 5-EU integrated signal density in naïve and GC B cells. Each symbol represents one cell. Scale bar 3μm. Representative of two independent experiments. H. 2D Airyscan ICC images of TFAM in naïve and GC B cells, with 3D reconstruction indicating individual mitochondrial nucleoids. Quantification of nucleoid number and volume. Each symbol represents one cell. Scale bar 1μm. Representative of two independent experiments. I. Deconvoluted confocal and STED super-resolution ICC images of TFAM in naïve and GC B cells. Scale bar 1μm. Representative of two imaging experiments Statistical significance was calculated by unpaired two-tailed t-test (B, G, H), Mann Whitney U test (F) or ordinary one-way ANOVA with Tukey’s multiple comparisons test (C, D)

To examine mitochondrial transcription, we next injected mice with 5-ethynyl uridine (5-EU), which is incorporated into actively synthesised RNA. We detected co-localisation between 5-EU and COX I, with 5-EU incorporation highly prominent in GC B cells, and most often seen in areas of high COX I expression (Fig. 1G).

To understand the regulation of this high rate of mitochondrial protein transcription and translation, we examined the expression and distribution of transcription factor A mitochondrial (TFAM). TFAM regulates transcription and translation of mitochondrial DNA, and is an effector of mitochondrial biogenesis. Using Airyscan immunocytochemistry in sorted GC B cells, we found TFAM protein organised into punctate structures representing mitochondrial nucleoids, which co-localised with 5-EU (Fig. 1G). These were more numerous and larger in GC B cells (Fig. 1H), and had a more elliptical morphology, which is associated with elevated transcriptional activity (Extended Data Fig. 1C)^26^. We confirmed the increase in nucleoid number using stimulated emission depletion (STED) super-resolution microscopy (Fig. 1I). Mitochondria are therefore highly dynamic in GC B cells, with prominent transcriptional activity associated with increased nucleoid content.

### 2. *Tfam* is essential for mitochondrial homeostasis during B cell development and differentiation

TFAM has been shown to have a role in the prevention of long term inflammatory immunosenescence in T cells, but to be dispensable for their development^20,27^. Whether TFAM is important for B cell development or function is unknown, as are the dynamics of ETC component expression in B cell developmental trajectories. To address these questions, we first conditionally deleted *Tfam* in B cells using *Cd79a*-Cre (hereafter B-Tfam), which is active from the early pro-B cell stage^28^. Tfam was efficiently deleted in mature splenic B cells (Extended Data Fig. 2A-B). B-Tfam mice appeared healthy, with no clinical signs of immunosenescence or overt autoimmunity. However, B-Tfam mice had a profound reduction in the peripheral B cell compartment (Fig. 2A, Extended Data Fig. 2C-E). Analysis of B cell development in the bone marrow indicated a failure of progression from the pro- to the pre-B cell stage, which represents an important physiological checkpoint in B cell ontogeny, and few cells were able to express surface immunoglobulin and become mature B cells (Fig. 2B-C, Extended Data Fig. 2F). We noted that heterozygous B-Tfam mice had normal B cell development and an intact peripheral B cell compartment (Extended Data Fig. 2G-H).

**Figure 2.**
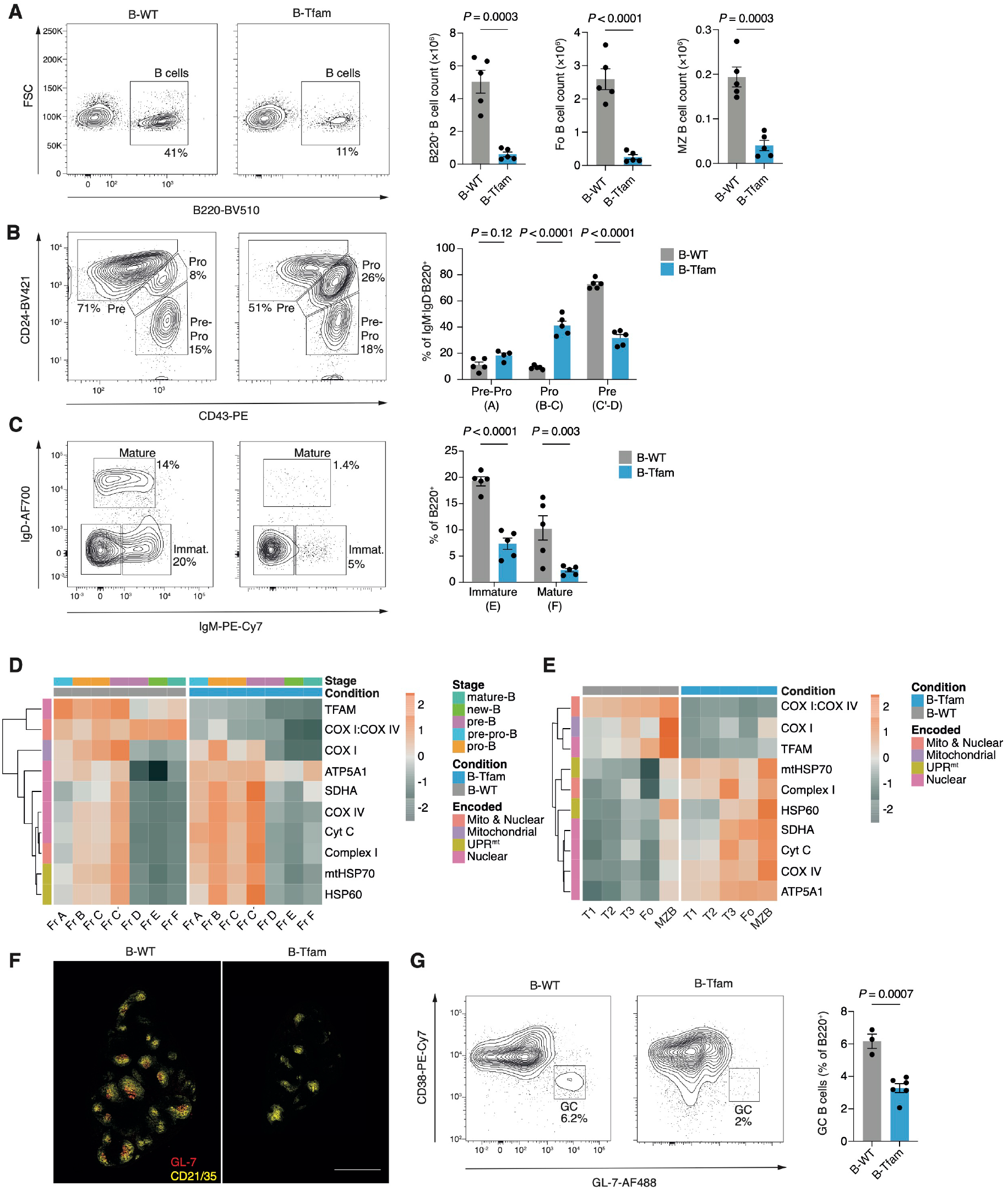
*Tfam* is essential for mitochondrial homeostasis during B cell development and differentiation. A. Representative flow cytometry plots of B220^+^ B cells from the spleens of unimmunised 6-week-old B-Tfam and B-WT mice, with quantification of total splenic B cell counts, and absolute counts of follicular (CD23^+^CD21^int^) and marginal zone (CD21^+^CD23^−^) B cell subsets (n=5). Representative of two independent experiments. B. Representative flow cytometry plots and quantification of bone marrow IgM^−^IgD^−^B220^+^ pre-pro, pro and pre-B cells isolated from B-Tfam and B-WT mice (n=5). Results representative of two independent experiments. Equivalent Hardy stages in parentheses. C. Representative flow cytometry plots and quantification of bone marrow-resident B220^+^ IgM^+^IgD^−^ immature and B220^+^IgD^+^ mature B cell subsets from B-WT and B-Tfam mice (n=5). Results representative of two independent experiments. Equivalent Hardy stages in parentheses. D. Heatmap of row Z-scores for gMFI of indicated mitochondrial proteins, measured by spectral flow cytometry in bone marrow B cell subsets (Hardy stages) of B-WT and B-Tfam mice (mean of two biological replicates). Results representative of two independent experiments with n=4 mice per group in total E. Heatmap of row Z-scores for gMFI of indicated mitochondrial proteins, measured by spectral flow cytometry in splenic B cell subsets of unimmunised B-WT and B-Tfam mice (mean of two biological replicates). Results representative of two independent experiments with n=3 mice per group in total. (F-G) B-WT and B-Tfam mice were immunised with SRBCs (day 0 & 5) intraperitoneally, and at day 12 spleens were analysed by immunohistochemistry (IHC) and flow cytometry. F. Representative tile-scan images of spleen sections from B-Tfam and B-WT mice depicting GL-7^+^ GCs and CD21/35^+^ B cell follicles. Scale bar 500μm. Results representative of two independent experiments G. Flow cytometry gating strategy and quantification of CD38^−^GL-7^+^ GC B cells in spleens of immunised B-Tfam (n=6) and B-WT mice (n=3). Results representative of two independent experiments. Statistical significance was calculated by unpaired two-tailed t-test (A, G), or two-way ANOVA with Šidák’s multiple comparison (B, C).

To understand the expression of ETC proteins during B cell development, and the effect of interference with mitochondrial gene transcription and translation by *Tfam* deletion, we quantified a panel of ETC proteins by high dimensional spectral flow cytometry in B-WT and B-Tfam bone marrow and spleen (Fig. 2D-E). We found that in B-WT bone marrow there was a progressive increase in the expression of most ETC proteins from the pre-pro B stage (Hardy fraction A^29^), peaking at the earliest pre-B subset (fraction C′), and then substantially falling in later stages (fractions D-F). Interestingly, TFAM expression peaked early (fraction A), implying that it might initiate a mitochondrial transcription program leading to upregulation of ETC proteins. The maximum expression of ETC proteins corresponded to the developmental block seen in B-Tfam mice (pro- to pre-B cell stage). B-Tfam mice deleted TFAM from fraction A onwards in bone marrow, and TFAM remained deleted in splenic B cell subsets, and accordingly mitochondrially-encoded COX I was downregulated. The nuclear-encoded ETC components COX IV, cytochrome C, ATP5A1, and succinate dehydrogenase A (SDHA) were increased.

In the periphery, ETC component expression was similar across transitional and follicular B cells, but increased in marginal zone B cells (Fig. 2E). TFAM deletion led to a much more marked upregulation of nuclear-encoded ETC proteins, which was maximal in marginal zone B cells. This was evident in a mismatch in the ratio of proteins encoded by nuclear or mitochondrial genomes in the same ETC complex – COX I and COX IV of complex IV. Seahorse extracellular flux analysis of unstimulated peripheral B cells showed no difference in oxygen consumption rate (OCR), but a significant increase in ECAR in B-Tfam mice (Extended Data Fig. 2I).

Also increased were the mitochondrial unfolded protein response (UPR^mt^)-associated proteins heat shock protein 60 (HSP60) and mitochondrial 70kDa heat shock protein (mtHSP70). These results demonstrate that ETC components are dynamically regulated across B cell developmental trajectories, and that TFAM is required for expression of mitochondrially-encoded proteins.

We next examined the ability of B-Tfam mice to generate GCs. We immunised B-Tfam mice with SRBCs and analysed GC formation by immunofluorescence and flow cytometry at day 12. Anatomical and immunofluorescence examination of B-Tfam spleens revealed smaller and fewer B cell follicles and an almost complete absence of GL-7^+^ GCs compared to B-WT mice (Fig. 2F), which was confirmed by flow cytometry (Fig. 2G). Despite the lack of developmental phenotype, *Tfam* heterozygosity in B cells led to a significantly reduced GC response following SRBC immunisation (Extended Data Fig. 2J). These results collectively suggest that TFAM is essential for normal B cell differentiation, and that in situations of low energy demand, partial respiratory compensation is possible despite substantially reduced translation of mitochondrial proteins.

### 3. GC B cells require TFAM

Having established a role for TFAM in B cell development and ETC complex balance, we next focused on its function in the GC reaction. To specifically delete *Tfam* in GC B cells, we generated Aicda-Cre × *Tfam* flox × Rosa26^STOP^tdTomato × *Prdm1*-mVenus mice (hereafter Aicda-Tfam), which allow simultaneous identification of cells which have expressed *Aicda* (activated, GC, memory B, and plasma cells) and/or currently express Blimp-1 (by detection of the fluorescent reporter proteins tdTomato and mVenus respectively) (Fig. 3A). Aicda-Tfam mice were immunised with SRBCs and analysed at day 12. Immunofluorescent staining of Aicda-Tfam spleen sections demonstrated small GCs in reduced numbers while B cell follicles, immunostained with anti-CD21/35, remained unaltered (Fig. 3B). Clusters of plasma cells (PCs) in the splenic cords were much smaller in Aicda-Tfam mice and Blimp1-mVenus^+^tdTomato+ cells were poorly represented, suggesting impairment of GC output (Fig. 3C). The proportion of CD138^+^ tdTomato^+^ long-lived plasma cells in bone marrow was also significantly lower (Extended Data Fig. 3A). We found that the generation of CD19^+^IgD^lo^CD38^−^GL-7^+^ (and correspondingly CD19^+^IgD^lo^CD38^−^tdTomato^+^) GC B cells was substantially reduced in Aicda-Tfam mice as measured by flow cytometry (Fig. 3D-E). We also confirmed a reduction in Blimp1-mVenus^+^tdTomato^+^ post-GC plasma cells (Fig. 3F).

**Figure 3.**
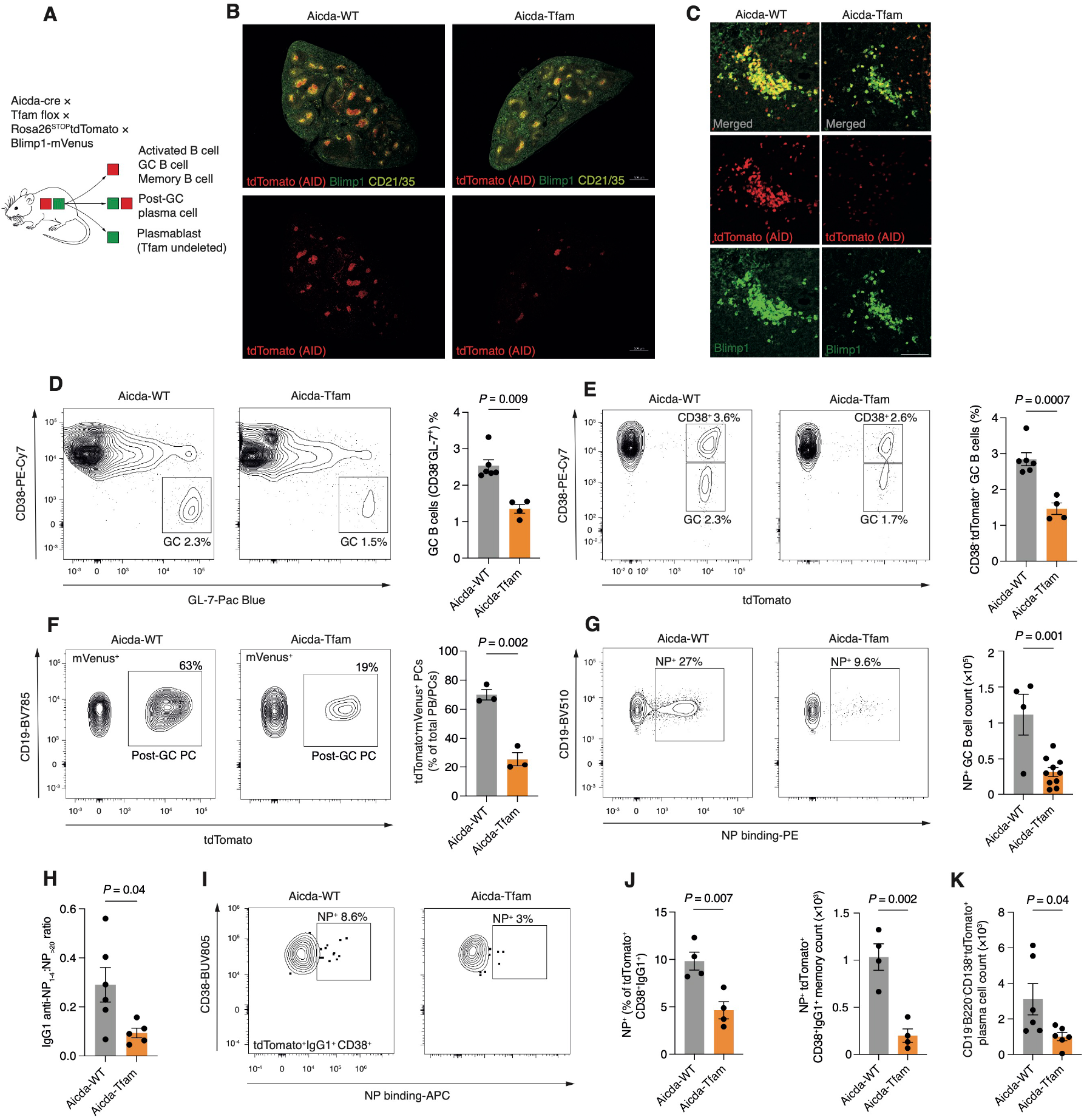
GC B cells require TFAM. A. Schematic of Aicda-Tfam mouse (Aicda-Cre × Tfam flox × Rosa26^STOP^tdTomato × Blimp1-mVenus) (B-F) Aicda-WT and Aicda-Tfam mice were immunised with SRBCs intraperitoneally, and on day 12 spleens were analysed by flow cytometry and/or immunohistochemistry (IHC) B. Representative Airyscan IHC confocal images of spleen sections from Aicda-Tfam and Aicda-WT mice. Scale bar 500μm. Representative of three independent experiments C. Airyscan images of plasma cell clusters in red pulp of spleen sections. tdTomato^+^Blimp1-mVenus^+^ double positive cells indicate post-GC plasma cells. Scale bar 75μm. D. Representative flow cytometry plots for the identification of GC B cells from Aicda-WT (n=6) and Aicda-Tfam (n=4) mice. Quantification of CD38^−^GL-7^+^ GC B cell percentages within the CD19^+^ B cell compartment. Representative of four independent experiments. E. Representative flow cytometry plots for the identification of GC B cells as CD38^−^ tdTomato^+^ in Aicda-WT (n=6) and Aicda-Tfam (n=4) mice. Results representative of four independent experiments. F. Representative flow cytometry plots depicting tdTomato^+^ Blimp1-mVenus^+^ post-GC plasma cells within the Dump^−^mVenus-Blimp1^+^ cell compartment. Comparison of tdTomato^+^ proportion within Dump^−^mVenus-Blimp1^+^ PB-PC compartment (n=3). Data representative of four independent experiments. (G-H) Aicda-WT and Aicda-Tfam mice were immunised with 50μg NP-CGG with Alum (at 1:1 ratio) intraperitoneally, and on day 14 spleens were harvested for flow cytometry and IHC analyses. G. Representative flow cytometry plots depicting CD19^+^CD38^−^GL-7^+^ GC B cells binding NP-PE or -APC from Aicda-Tfam (n=4) or Aicda-WT (n=10). Quantification of NP-specific GC B cell absolute counts. Data pooled from and representative of three independent experiments. H. ELISA quantification and comparison of the ratio of IgG1 NP-specific high-affinity antibodies to low-affinity antibodies detected by binding to NP1-4 and NP>20 antigens respectively from Aicda-WT (n=6) and Aicda-Tfam (n=5). Data pooled from two independent experiments. (I-K) Aicda-WT and Aicda-Tfam mice were immunised with 50μg NP-CGG with Alum (at 1:1 ratio) intraperitoneally, and on day 30 boosted with 50μg NP-CGG in PBS. Spleens and bone marrow were harvested for flow cytometry at day 70. I. Representative flow cytometry plots depicting tdTomato^+^ IgG1^+^ CD38^+^ memory B cells binding NP-APC from Aicda-WT and Aicda-Tfam mice. J. Proportional comparison and absolute count quantification of NP-binding within CD38^+^ IgG1^+^ tdTomato^+^ memory B cells from Aicda-WT and Aicda-Tfam mice (n=4). Representative of two independent experiments. K. Absolute plasma cell counts in bone marrow of Aicda-WT and Aicda-Tfam (n=6) at day 70. Data pooled from two independent experiments. Statistical significance was calculated by unpaired two-tailed t-test.

Next, we examined antigen-specific GC formation by immunising mice with 4-hydroxy-3-nitrophenylacetyl-chicken gamma globulin (NP-CGG). After 14 days, the proportion and absolute numbers of NP-binding GC B cells were much reduced in Aicda-Tfam mice, as were those of PCs measured in situ, and the PC to GC B cell ratio was correspondingly decreased (Fig. 3G and Extended Data Fig. 3B-D). Affinity maturation was also compromised, with decreased binding of IgG1 antibodies to NP_1-4_ compared with NP_>20_ (Fig. 3H and Extended Data Fig. 3E-F). There was no difference in IgM anti-NP antibodies however, in keeping with the extrafollicular origin of this response, produced by plasmablasts which have not expressed *Aicda* (Extended Data Fig. 3G).

To understand if loss of *Tfam* compromised B cell memory, we immunised Aicda-Tfam and Aicda-WT mice with NP-CGG in alum and then boosted them at D30 with NP-CGG in PBS. At D70 there were substantially fewer NP-binding CD38^+^IgG1^+^ memory B cells (Fig. 3I-J) and at day 49 a reduction in IgG1 anti-NP antibodies reactive against NP_1-4_ and NP_>20_ (Extended Data Fig. 3H). The number of plasma cells in the bone marrow was also significantly reduced at day 70 (Fig. 3K).

Mitochondria play a central role in the regulation of apoptosis, but surprisingly the apoptosis rate detected by means of activated caspase 3 staining in Aicda-Tfam GC B cells was comparable with that of Aicda-WT controls (Extended Data Fig. 3I). In situ terminal deoxynucleotidyl transferase dUTP nick end labelling (TUNEL) also demonstrated unaltered apoptosis (Extended Data Fig. 3J). We next evaluated the cell cycle dynamics of Aicda-Tfam GC B cells, first assessing G2 and M stages by measuring cyclin B1 and phospho-H3 (p-H3) expression, which were not significantly different between Aicda-Tfam and Aicda-WT mice (Extended Data Fig. 3K). 5-ethynl-2-deoxyurine (EdU) incorporation into DNA, which detects S phase, was also unaffected (Extended Data Fig. 3L).

We also confirmed that *Tfam* deletion did not affect proliferation or cell viability in vitro, by treating B cells from *Tfam* flox × Rosa26^STOP^tdTomato or wild type Rosa26^STOP^tdTomato mice with TAT-Cre recombinase, then stimulating them with anti-IgM, agonistic anti-CD40, and IL-4 for four days (Extended Data Fig 3M). *Tfam* was effectively deleted in tdTomato^+^ cells (Extended Data Fig 3N), but there was no difference in dilution of CellTrace Violet or in viability (Extended Data Fig 3M-O).

These data demonstrate that loss of *Tfam* markedly compromises GC B cell differentiation and output, but without detectable effects on proliferation and survival in those cells already committed to GC fate.

### 4. TFAM controls transcriptional entry into the GC program

In order to examine the effects of *Tfam* deletion on the transcriptional program of GC B cells, we performed combined single cell gene expression profiling and V(D)J sequencing. We immunised Aicda-Tfam and Aicda-WT mice with NP-CGG, and sorted tdTomato^+^ cells at day 14 (Fig. 4A). This population includes any B cell that has expressed *Aicda*, and so will include pre-GC, GC, memory, and plasma cells. We sequenced a total of 9,948 cells from Aicda-Tfam and 9,667 cells from Aicda-WT mice. Cells were clustered following integration of the two experimental groups, and clusters were identified by canonical markers (Fig. 4B-C). We identified 9 shared clusters, broadly separated into B cells from GC, CD38^+^ non-GC, and plasma cell populations. Cluster 0 (C0), which expressed markers of immaturity suggestive of an activated precursor (AP) state (*Ighd*, *Ccr6*, *Gpr183*, *Cd38*, *Sell*, and low levels of *Bcl6*), was significantly expanded in Aicda-Tfam compared with Aicda-WT mice (Fig. 4D).

**Figure 4.**
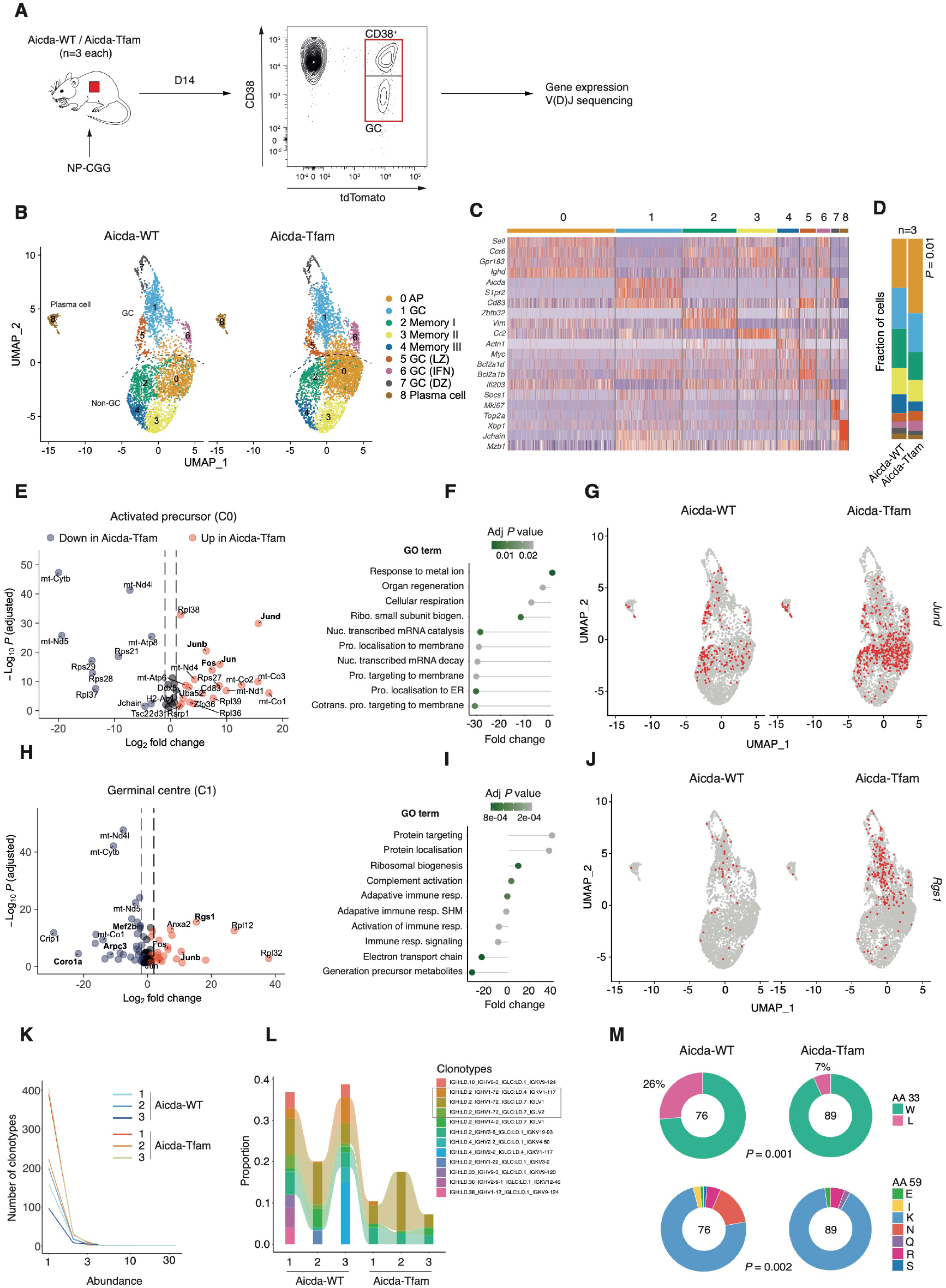
TFAM controls transcriptional entry into the GC program. A. Schematic of experimental design. Aicda-WT and Aicda-Tfam mice (n=3 per group) were immunised with 50μg NP-CGG with Alum (at 1:1 ratio) intraperitoneally and live Dump^−^ (CD3^−^, Gr1^−^, CD11c^−^) tdTomato^+^ cells were flow sorted before bead capture and 10x library preparation and sequencing. B. UMAP projections and clustering of integrated Aicda-WT and Aicda-Tfam datasets. C. Heatmap of selected differentially expressed marker genes by cluster for all samples. D. Cluster proportions between groups. E. Volcano plot of differentially expressed genes in cluster 0 (activated precursors). F. Single cell pathway analysis (SCPA) of Gene Ontogeny Biologicals Processes (GO BP) in AP cluster. G. *Jund* gene expression projected onto clustered data. H. Volcano plot of differentially expressed genes in cluster 1 (germinal centre). I. SCPA of GO BP in germinal centre cluster (cluster 1). J. *Rgs1* gene expression projected onto clustered data. K. Plot of clonotype abundance distribution for each sample. L. Plot of proportions of top clonotypes for each sample. Clonotypes with *Ighv1-72* usage are highlighted. M. Quantification of substitution at *Ighv1-72* amino acid coding positions 33 and 59 for GC B cell cluster. Number of GC B cells with *Ighv1-72* usage analysed indicated on chart. Statistical significance was calculated by t-test with correction for multiple comparison by the Bonferroni methods (D, E, H), or Chi-square test (M)

Examination of differential gene expression in C0 revealed, as expected, broad dysregulation of mitochondrial gene expression in Aicda-Tfam mice, in keeping with the function of TFAM as a regulator of mitochondrial transcription (Fig. 4E). Pathway analysis comparing multivariate distributions (SCPA)^30^ demonstrated substantial downregulation of translation initiation and elongation gene sets (Fig. 4F). Gene components of the AP-1 signalling pathway *Jun*, *Junb*, and *Fos* were all significantly upregulated (Fig. 4G). The AP-1 pathway is broadly upregulated by cellular stress signalling, reactive oxygen species, and in response to environmental cues^31,32^.

Analysis of the main GC cluster (C1) demonstrated as before, dysregulation of mitochondrial and ribosomal gene transcription, with upregulation of *Jun, Fos*, and *Junb* (figure 4H-I). Notably increased in Aicda-Tfam cells was *Rgs1* (regulator of G-protein signaling-1), a GTPase activating protein which has an important role in the negative regulation of cell movement in response to chemokines (e.g. CXCL12)^33,34^ (Fig. 4J). Downregulated was *Coro1a* (coronin-1), which encodes an actin-binding protein required for cell migration^35^, and *Arpc3* (actin-related protein 2/3 complex subunit 3), which mediates branched actin polymerisation and actin foci formation^36^. We also detected a reduction in *Mef2b*, which regulates GC enhancer genes by transactivating *Bcl6*, and is important for GC B cell positioning^37^.

Across other clusters the dysregulation of mitochondrial and ribosomal genes in Aicda-Tfam cells was repeated. Cluster 6 within the GC supercluster was characterised by expression of interferon response genes, and was subjectively expanded in Aicda-Tfam mice, although this did not reach statistical significance. This may represent a response to mitochondrial DNA leaking into the cytoplasm, which has been previously noted in Tfam deficient cells^21^.

We next examined B cell clonality by evaluation of V(D)J sequences. Since we sorted all tdTomato^+^ B cells, the overall clonal diversity of all samples remained high (Fig. 4K). As expected in the anti-NP immune response, V_H_1-72 gene usage was dominant (Fig. 4L).

However, there was more diversity in Aicda-Tfam mice, with fewer larger clones, suggesting that there was less ability for evolution of dominant clones. There was significantly less somatic hypermutation in Aicda-Tfam B cells, including within the *Ighv1-72* gene in GC B cells (Extended Data Fig. 4A-C). The W33L substitution in CDR1, and substitutions of K59 of the V_H_1-72 heavy chain confer increased affinity for NP^38,39^. There were significantly lower W33L and K59 substitution rates in the Aicda-Tfam GC B cell cluster, in keeping with our observation that high-affinity NP binding was reduced (Fig. 4M). There was no W33L substitution and negligible K59 mutation in the AP cluster (C0) of either Aicda-WT or Aicda-Tfam mice, reflecting their pre-GC state (Extended Data Fig. 4D).

### 5. TFAM is required for GC B cell commitment

Given the relative accumulation in immunised Aicda-Tfam mice of activated precursor B cells with high expression of *Sell* (L-selectin), *Ccr6*, and a GC transcriptional profile suggestive of altered cell trafficking and cytoskeleton dynamics, we hypothesised that *Tfam* is required for activated B cells to enter the GC and remain appropriately spatially positioned.

We first confirmed the proportional expansion of APs, defined as tdTomato^+^CD38^+^IgD^+^, in Aicda-Tfam mice following NP-CGG immunisation (Fig. 5A). Despite a significant numerical reduction in GC B cells, the AP population was maintained (Extended Data Fig 5A). This was also seen to a more striking extent in B-Tfam mice immunised with SRBC, with APs defined as IgD^+^GL-7^int^ (Extended Data Fig. 5B). We found that in Aicda-WT mice, a lower proportion of AP B cells bound NP compared to GC B cells, and did so with lower affinity, in keeping with their lack of W33L mutation and low levels of somatic hypermutation (Extended Data Fig. 5C).

**Figure 5.**
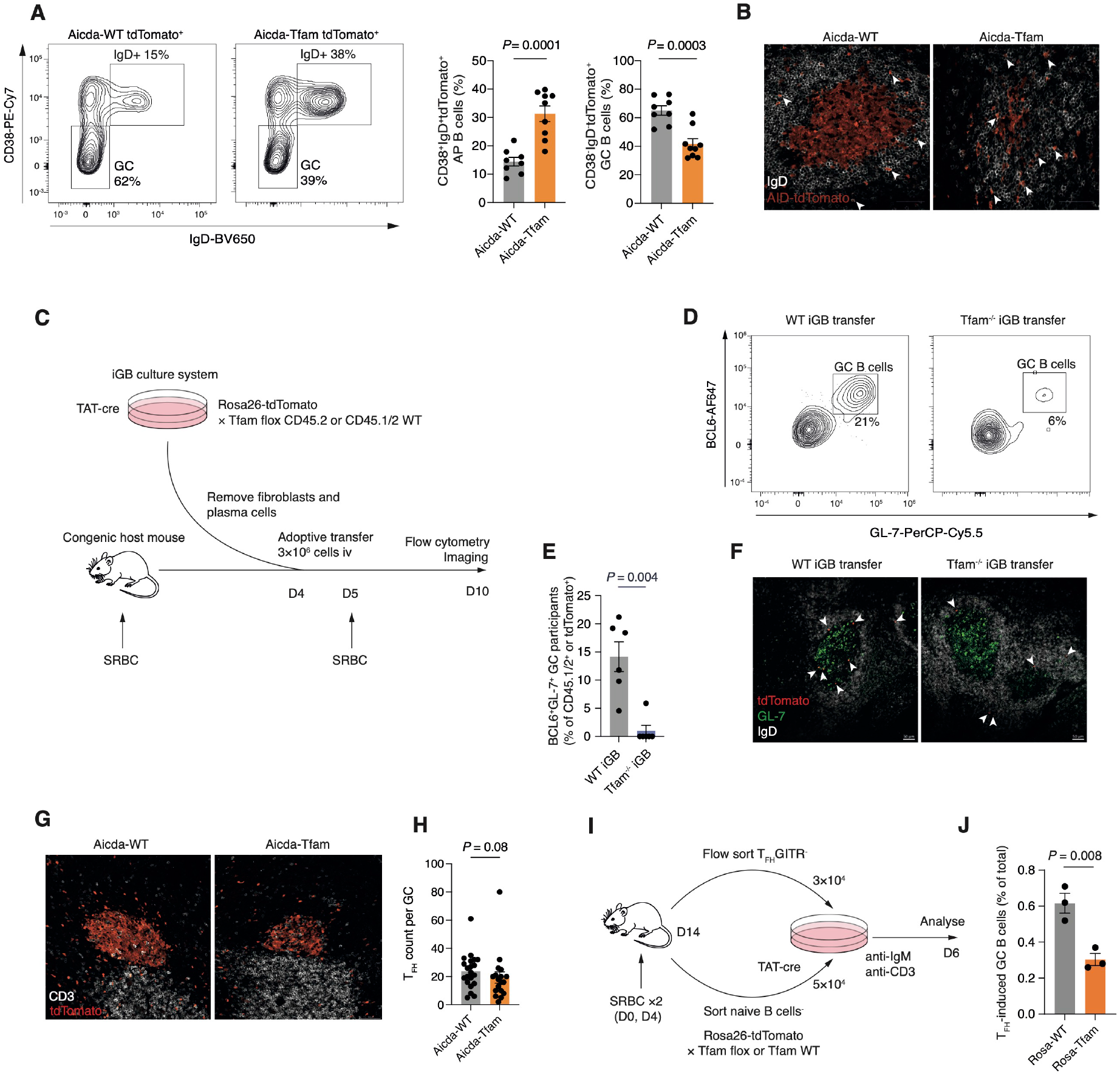
TFAM is required for GC B cell commitment. A. Representative flow cytometry plots depicting gating of CD38^+^IgD^+^ activated precursors (AP) and CD38^−^IgD^−^ GC B cells within the CD19^+^tdTomato^+^ population. Quantification of AP and GC B cell percentages from NP-CGG immunised Aicda-WT (n=8) and Aicda-Tfam mice (n=9). Data pooled from two independent experiments and representative of four independent experiments. B. Representative IHC images of GCs from Aicda-WT and Aicda-Tfam mice. Arrows indicate tdTomato^+^IgD^+^ APs. 20× magnification, scale bar 20μm. Representative of two independent experiments. C. Schematic of experimental design for iGB culture and adoptive transfer to assess in vivo GC entry/recruitment. D. Flow cytometric characterisation of GC entry of adoptively transferred WT and Tfam^−/−^ iGB cells in SRBC-immunised CD45.1 congenic hosts. GC B cells defined as (BCL6^+^ GL-7^+^). Representative of two independent experiments. E. Quantification of D (n=6) F. Confocal images of splenic sections depicting tdTomato^+^ WT and Tfam^−/−^ iGB cells adoptively transferred as in C into separate congenic hosts. Arrows indicate transferred tdTomato^+^ iGB cells. Representative of two independent experiments. G. Representative confocal images of the splenic T_FH_ cell compartment in Aicda-WT and Aicda-Tfam mice. Scale bar 50μm. Representative of three independent experiments. H. T_FH_ cell counts per GC from immunised Aicda-WT and Aicda-Tfam mice. Each symbol represents one GC pooled from n=4 mice. Representative of three independent experiments. I. Schematic of experimental design of T_FH_-B co-culture. CD4^+^CD19^−^CXCR5^+^ICOS^+^GITR^−^ T_FH_ cells (3×10^4^) were sorted from wild type mice immunised with SRBC (at days 0 and 4 for enhanced GC reaction) and co-cultured with TAT-Cre treated naïve B cells (5×10^4^) from Aicda-WT and Aicda-Tfam mice, in the presence of anti-CD3 and anti-IgM for six days. J. Percentages of in vitro T_FH_-induced Aicda-WT or Aicda-Tfam GC B cells (GL-7^+^ tdTomato^+^ IgG1^+^) (of total viable cells). Technical replicates of n=2 pooled biological replicates shown. Representative of two independent experiments with n=3 biological replicates in total. Statistical significance was calculated by unpaired two tailed t-test.

We then examined splenic sections of immunised Aicda-Tfam mice under high magnification (Fig. 5B). We found highly disorganised GC architecture, with poor GC B cell compartmentalisation, and within the follicle, there were relatively many more tdTomato^+^IgD^+^ B cells in Aicda-Tfam mice. The level of BCL6 protein, a master transcriptional regulator of GC commitment and entry was lower in Aicda-Tfam GC B cells (Extended Data Fig. 5E), in keeping with the reduced transcription of its activator *Mef2b* we observed in our single cell gene expression data.

We next asked whether it was possible to overcome the failure of AP B cells to enter the GC by adoptively transferring pre-activated Tfam^−/−^ AP-like B cells into primed CD45.1 mice. We used the iGB B cell culture system^40^ and TAT-Cre to delete *Tfam* in *Tfam* flox × Rosa26-stop-tdTomato B cells, or with wild type congenically-marked Rosa26^STOP^tdTomato × CD45.1/2 control B cells. This experimental design allows competitive transfer of activated B cells to take place (Fig. 5C). The resulting Tfam^−/−^ iGB cells effectively deleted TFAM at day 4 of culture, and this resulted in decreased expression of mitochondrially-encoded ETC proteins and upregulation of nuclear-encoded proteins, as seen during B cell development in B-Tfam mice (Extended Data Fig. 5F). Loss of Tfam did not affect cell expansion over four days of culture (Extended Data Fig. 5G). However, Tfam^−/−^ iGB cells were at a striking competitive disadvantage in GC participation following adoptive transfer (Fig. 5D-E, Extended Data Fig. 5H), confirmed by immunofluorescent imaging (Fig. 5F).

Activated B cells need help from cognate T_FH_ cells in order to enter the GC, and in turn T_FH_ cells require interaction with activated B cells to develop and survive^41^. We reasoned that the defects we observed upon deletion of *Tfam* might be due to abnormalities in T_FH_-B cell interaction, or a defective T_FH_ pool. However, following immunisation, we did not detect a numerical defect in the T_FH_ compartment (Fig. 5G-H). We then co-cultured sorted ex vivo T_FH_ cells (for gating strategy see Extended Data Fig. 5D) with TAT-Cre-treated Rosa26^STOP^tdTomato × Tfam flox or wild type control B cells (Fig. 5I)^42^. We noted a reduction in tdTomato^+^GL-7^+^IgG1^+^ induced GC (iGC) B cells following *Tfam* deletion (Fig 5J). The capacity of iGB cells to process antigen and form conjugates with T cells in vitro was intact (Extended Data Fig. 5I). To determine whether the defect we saw in Aicda-Tfam mice was B cell-intrinsic, we generated mixed competitive bone marrow chimeras with CD45.1 congenic wild type mice, in which T_FH_ generation would be intact. Aicda-Tfam GC B cells were outcompeted by CD45.1 wild type cells, compared with Aicda-WT controls in spleens and Peyer’s patches (Extended Data Fig. 5J). Overall these data suggest that Tfam is required for entry of activated B cells into the GC, and that this defect is principally cell-intrinsic.

### 6. TFAM regulates mitochondrial translation and metabolic homeostasis in activated B cells

We next examined the expression of TFAM and mitochondrial ETC components following immunisation. There was dynamic expression of TFAM, which peaked at the AP stage, and then subsided following GC entry, and a progressive increase in COX I, maximal in GC B cells (Fig. 6A-C). Other ETC proteins were also highly expressed in GC B cells, including those encoded in nuclear and mitochondrial DNA. Deletion of *Tfam* led to a program of alteration in mitochondrial ETC protein expression in AP and GC B cells (Fig. 6C), much like that observed during B cell development.

**Figure 6.**
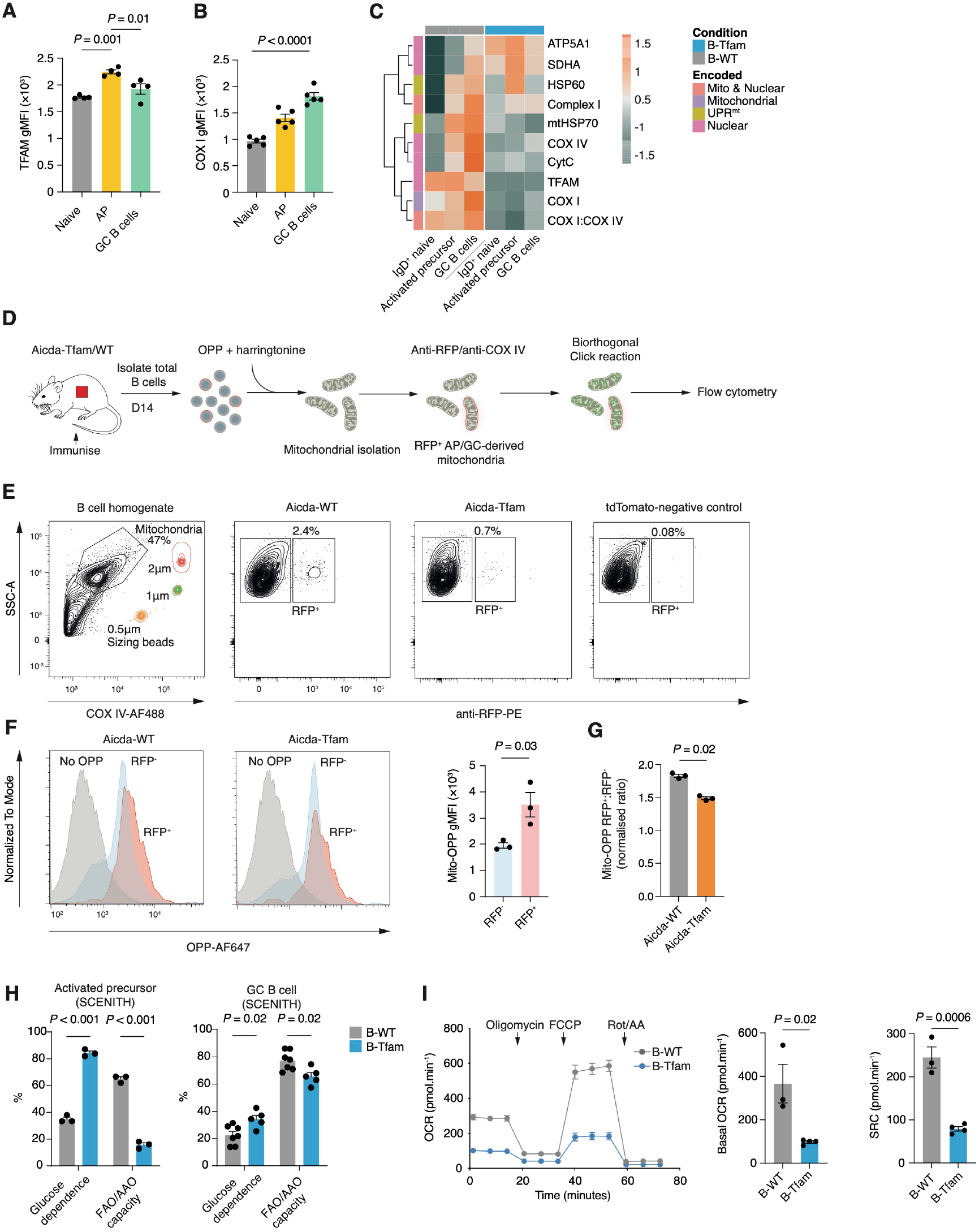
TFAM regulates mitochondrial translation and metabolism in activated B cells. A. Quantification of TFAM gMFI in tdTomato^+^CD38^−^ GC B cells, tdTomato^+^IgD^+^ AP, and tdTomato^−^IgD^+^ naïve B cell compartments from SRBC immunised Aicda-WT and Aicda-Tfam mice (n=5). Data representative of three independent experiments. B. Quantification of COX I gMFI in tdTomato^+^CD38^−^ GC B cells, tdTomato^+^IgD^+^ AP, and tdTomato^−^IgD^+^ naïve B cell compartments from SRBC immunised Aicda-WT and Aicda-Tfam mice (n=5). Data representative of three independent experiments. C. Heatmap of row Z-scores for gMFI of indicated mitochondrial proteins, measured by spectral flow cytometry in splenic B cell subsets of NP-CGG immunised (D14) B-WT and B-Tfam mice (mean of 3 biological replicates). Results representative of two independent experiments with n=4 mice per group in total. D. Schematic of single mitochondrion translation assay E. Gating strategy for isolated mitochondria based on side scatter-A (SSC-A), sizing beads, and COX IV-AF488 expression, and for RFP^+^ AP-GC-derived mitochondria F. FACS histogram plots depicting OPP-AF647 signal in AP-GC derived RFP^+^ and naïve B cell-derived RFP^−^ mitochondria from immunised Aicda-Tfam and Aicda-WT mice. Quantification of OPP gMFI in RFP^+^ vs RFP^−^ mitochondria from immunised Aicda-WT (n=3). Data pooled from three independent experiments. G. Normalised OPP in RFP^+^ AP-GC-derived mitochondria from Aicda-Tfam and Aicda-WT mice. The experiment was performed in three batches and values were pooled after batch correction. H. Quantification of glucose dependence and fatty acid oxidation (FAO) or amino acid oxidation (AAO) capacity of cells based on OPP incorporation in splenic IgD^+^GL-7^int^ AP and IgD^−^GL-7^+^CD38^−^ GC B cells from B-WT and B-Tfam mice (n=3-5), treated ex vivo with metabolic inhibitors (oligomycin and/or 2-DG). Data pooled from two independent experiments. See methods for details of SCENITH calculations. I. Real-time OCR measurements (from MitoStress test) of 3×10^5^ B cells stimulated overnight (anti-CD40 and IL-4) from B-Tfam (n=4) and B-WT (n=3) mice, with quantification of basal OCR and spare respiratory capacity (SRC). Results pooled from two independent experiments. Statistical significance was calculated by one-way ANOVA with Dunnett’s multiple comparison test (A-B), unpaired two tailed t-test (F, I), unpaired two tailed t-test following batch correction (G) or two-way ANOVA with Šidák’s multiple comparison (H).

To directly confirm the activity of mitochondrial translation, we then measured incorporation of the amino acid analogue O-propargyl-puromycin (OPP) in isolated ex vivo mitochondria from Aicda-WT and Aicda-Tfam B cells by flow cytometry (Fig. 6D). We first treated isolated total splenic B cells from immunised mice with OPP, and harringtonine, which blocks ribosomal protein synthesis in the cytoplasm, but does not affect mitochondrial ribosomes^43^. As tdTomato is ubiquitously distributed and enters mitochondria^44^ (Extended Data Fig. 6A), we were able to use it as a marker of *Aicda* expression and therefore AP- or GC B cell-derived origin following cellular homogenisation. We detected OPP using Click chemistry, and immunostained for COX IV and tdTomato (with anti-RFP). Mitochondria were identified by flow cytometry using submicron sizing beads, COX IV expression, and side scatter properties (Fig. 6E). Within the mitochondrial pool, an RFP^+^ population was readily identified. The proportion of RFP^+^ mitochondria was substantially lower in Aicda-Tfam mice, reflecting their defective GC formation (Fig. 6E). We detected significantly more OPP uptake in RFP^+^ mitochondria compared to those of non-AP or GC B cell origin (RFP^−^)(Fig. 6F). Translation was decreased in RFP^+^ mitochondria from Aicda-Tfam B cells compared to Aicda-WT (Fig. 6G), and using Tfam^−/−^ iGB cells, treatment with the mitochondrial translation inhibitor and oxazolidinone antibiotic chloramphenicol^45^ did not lead to additional suppression (Extended Data Fig. 6B).

We then examined the metabolic consequences of Tfam deletion on APs and GC B cells, quantifying protein translation rate by the incorporation of OPP as a proxy for metabolic capacity, as recently reported^46^. The addition of inhibitors of glycolysis (2-deoxyglucose) or ATP synthetase (oligomycin) which disrupts the electron transport chain allows the relative contributions of these metabolic pathways to be estimated. GC B cells had significantly higher basal OPP incorporation than AP B cells, reflecting their high levels of metabolic activity (Extended Data Fig. 6C). Loss of *Tfam* led to an increase in glucose dependence and a marked reduction in fatty acid oxidation capacity in AP B cells, seen to a lesser extent in GC B cells (Fig. 6H and Extended Data Fig. 6C-D. These results were mirrored by Seahorse extracellular flux analysis in B cells stimulated overnight with agonistic anti-CD40 and IL-4, and iGB cells, which demonstrated a substantially reduced OCR and spare respiratory capacity (SRC) following *Tfam* deletion (Fig. 6I and Extended Data Fig. 6E-F).

### 7. Tfam deletion disrupts GC spatial organisation and impairs cell mobility

The germinal centre is a spatially organised structure divided into light and dark zones. We found that Aicda-Tfam GCs were poorly compartmentalised, with smaller DZs on immunohistochemistry (Fig. 7A), and a disrupted DZ/LZ ratio measured using flow cytometry (Fig. 7B).

**Figure 7:**
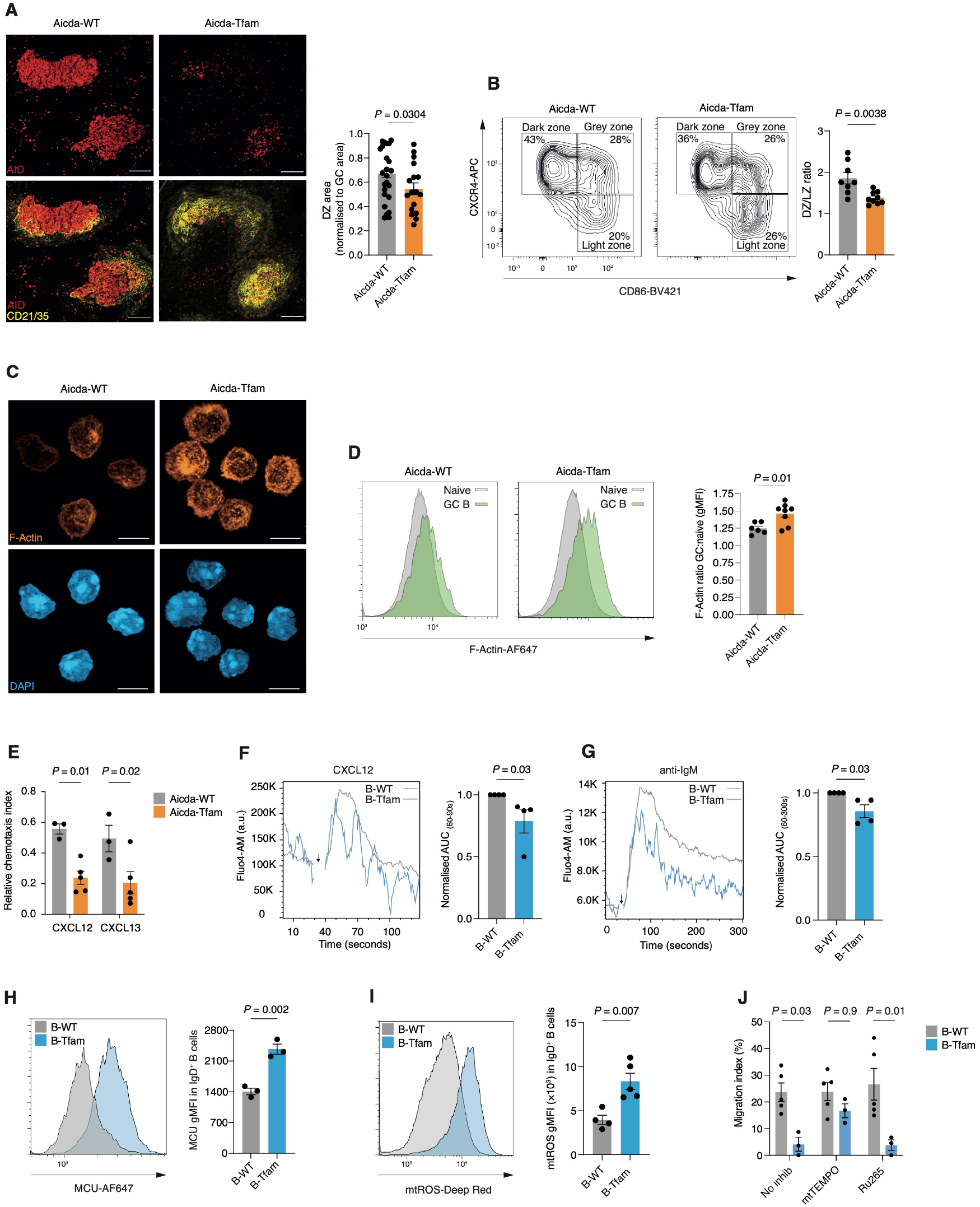
*Tfam* deletion disrupts GC spatial organisation and impairs cell mobility. A. Representative Airyscan IHC images of spleen sections from SRBC-immunised Aicda-WT and Aicda-Tfam mice. Quantification of DZ area normalised to GC area. Each symbol represents an individual GC. Image analyses were performed including all GCs identified on two sections from 2 mice of each genotype. Scale bar 100μm. Representative of >3 independent experiments. B. Representative flow cytometry plots of DZ, GZ, and LZ subpopulations of tdTomato^+^CD38^−^GL-7^+^ GC B cells, based on CXCR4 and CD86 expression. Quantification of the DZ to LZ ratio from NP-CGG immunised Aicda-WT (n=8) and Aicda-Tfam mice (n=9). Representative of four independent experiments. C. 3D Airyscan confocal image of magnetic bead-sorted and F-actin phalloidin + DAPI-stained GC B cells from Aicda-WT and Aicda-Tfam mice. All cells imaged expressed tdTomato. Scale bar 6μm. Representative of two independent experiments D. Representative flow cytometry histogram of F-actin phalloidin fluorescence of tdTomato^+^GL-7^+^ GC B cells and tdTomato^−^IgD^+^ naïve B cells from immunised Aicda-WT (n=6) and Aicda-Tfam mice (n=8). Quantification of F-actin phalloidin gMFI in tdTomato^+^IgD^−^GL-7^+^ GC B cells from immunised Aicda-WT and Aicda Tfam mice normalised to tdTomato^−^IgD^+^ naïve B cells from the same host. Data pooled from three independent experiments. E. Quantification of chemotaxis to CXCL12 and CXCL13 in GC B cells from Aicda-WT and Aicda-Tfam mice (n=4). Data representative of four independent experiments. F. Representative Fluo4-AM geometric mean signal intensity kinetics (moving average) of B-WT and B-Tfam B220^+^ B cells stimulated with CXCL12 (200ng/ml) for 120s. Quantification of area under curve (AUC) between 60-90s (corresponding to peak response). Experiment was run as technical duplicates in four independent replicate experiments consisting of one B-Tfam and one wild-type mouse. Data points from B-Tfam were normalised to wild type data run in the same batch. G. Representative Fluo4-AM gMFI signal kinetics (moving average) of B-WT and B-Tfam B220^+^ B cells stimulated with anti-IgM (10μg/ml) for 300s. Quantification of AUC between 60-300s. Experiment was run as technical duplicates in four independent replicate experiments consisting of one B-Tfam and one wild-type mouse in each. Data points from B-Tfam were normalised to wild-type data run in the same batch. H. Representative flow cytometry histogram of MCU fluorescence in B220^+^ IgD^+^ B cells from unimmunised B-Tfam and B-WT mice. Quantification of MCU gMFI in IgD^+^ B cells from B-Tfam and B-WT mice (n=3). Data representative of three independent experiments. I. Representative flow cytometry histogram of mtROS Deep Red fluorescence in IgD^+^ B cells from B-WT (n=4) and B-Tfam mice (n=5). Data representative of two independent experiments. J. Quantification of chemotaxis to CXCL12 (100ng/ml) in naive B cells from B-Tfam (n=3) and B-WT mice (n=5) in the presence of mtSOX scavenger mitoTEMPO (100μM) or MCU inhibitor Ru265 (30μM). Data representative of two independent experiments and pooled after batch correction. Statistical significance was calculated by unpaired two-tailed t-test (B,D,I), Mann Whitney U (F,G) or two-way ANOVA with Šidák’s multiple comparison test (E), with batch as a covariate (J).

Positioning of B cells in the GC is controlled by the chemokines CXCL12 and CXCL13, which promote migration to the DZ and LZ, respectively^47^. Since our preceding data indicated that AP B cells need TFAM to enter the GC, and given the transcriptional profile of Aicda-Tfam GC B cells was suggestive of cytoskeletal and mobility defects, we hypothesised that TFAM was required for proper cellular positioning in GCs.

We examined the cellular actin network of TFAM-deficient B cells, and found a significant increase in filamentous actin (F-actin) in Aicda-Tfam B cells (Fig. 7C-D), which was also evident in B-Tfam B cells (Extended Data Fig. 7A-B). Rearrangement of the actin cytoskeleton is critical for B cell migration^48^, and to understand if Tfam deletion compromised GC B cell motility, we performed a transwell migration assay to determine chemotaxis in response to the chemokines CXCL12 and CXCL13. We found that Aicda-Tfam GC B cells migrated poorly, with significantly reduced chemotaxis compared to Aicda-WT cells (Fig. 7E).

B-cell receptor (BCR) and chemokine-driven cytoplasmic calcium mobilisation critically regulates F-actin organisation in B cells^49^. As mitochondria are an important reservoir of intracellular calcium^50^, we asked whether Tfam deletion led to dysregulated intracellular calcium levels. Following CXCL12 stimulation, B cells from B-Tfam mice failed to sustain peak cytoplasmic calcium levels (Fig. 7F), which we also observed after BCR stimulation with anti-IgM antibodies (Fig. 7G). Interestingly, we also detected a significant upregulation in levels of the mitochondrial calcium uniporter (MCU), suggesting elevated mitochondrial calcium uptake capacity in B-Tfam B cells, whereas CD3^+^ T cells showed comparable MCU expression (Figure 7H & Extended Data Fig. 7C).

Mitochondrial ROS were also substantially increased in B-Tfam naïve and APs (Fig. 7I and Extended Data Fig. 7D). ROS activate the AP-1 signalling pathway^32^, and this observation was therefore consistent with the transcriptional profile of Aicda-Tfam AP B cells.

Given these observations, we next tested whether either suppression of mitochondrial ROS with the scavenger mitoTEMPO^51^, or inhibition of MCU function with the ruthenium compound Ru265^52^ improved cell motility in B-Tfam B cells in response to CXCL12. MCU inhibition did not improve transwell migration, but strikingly mitoTEMPO largely rescued the defect in B-Tfam cells (Fig. 7J).

Our data therefore collectively suggest that TFAM is required for proper cellular motility to enable entry into and movement within the GC, and that this effect is associated with ETC dysfunction and increased mitochondrial ROS.

### 8. *Tfam* deletion in B cells abrogates the development and progression of lymphoma

Having demonstrated an essential role for TFAM in B cell development and entry into the GC, we next asked whether it was required for the development of lymphoma. One of the most common mutations giving rise to DLBCL is translocation of *MYC* to immunoglobulin gene loci, leading to its unregulated expression. c-Myc is a key transcription factor regulating cell cycle and growth, cellular metabolism, and mitochondrial biogenesis^53^. We reasoned that deletion of *Tfam* would counter the oncogenic effects of c-Myc overexpression. We employed the well-established Eμ-Myc transgenic mouse model of lymphoma, in which *Myc* is expressed under the control of the *Igh* enhancer^54^. Eμ-Myc mice develop lymphoma with high penetrance from a median age of 11 weeks. To understand what effects overexpression of Myc had on mitochondrial translation, we transferred established lymphoma cells from Eμ-Myc mice (CD45.2) into wild type CD45.1 congenic hosts, to allow us to compare B cells from the same environment (Fig. 8A). We found very markedly higher expression of COX I in CD45.2 lymphoma cells compared with wild type B cells, with significantly upregulated TFAM (Fig. 8B-C).

**Figure 8:**
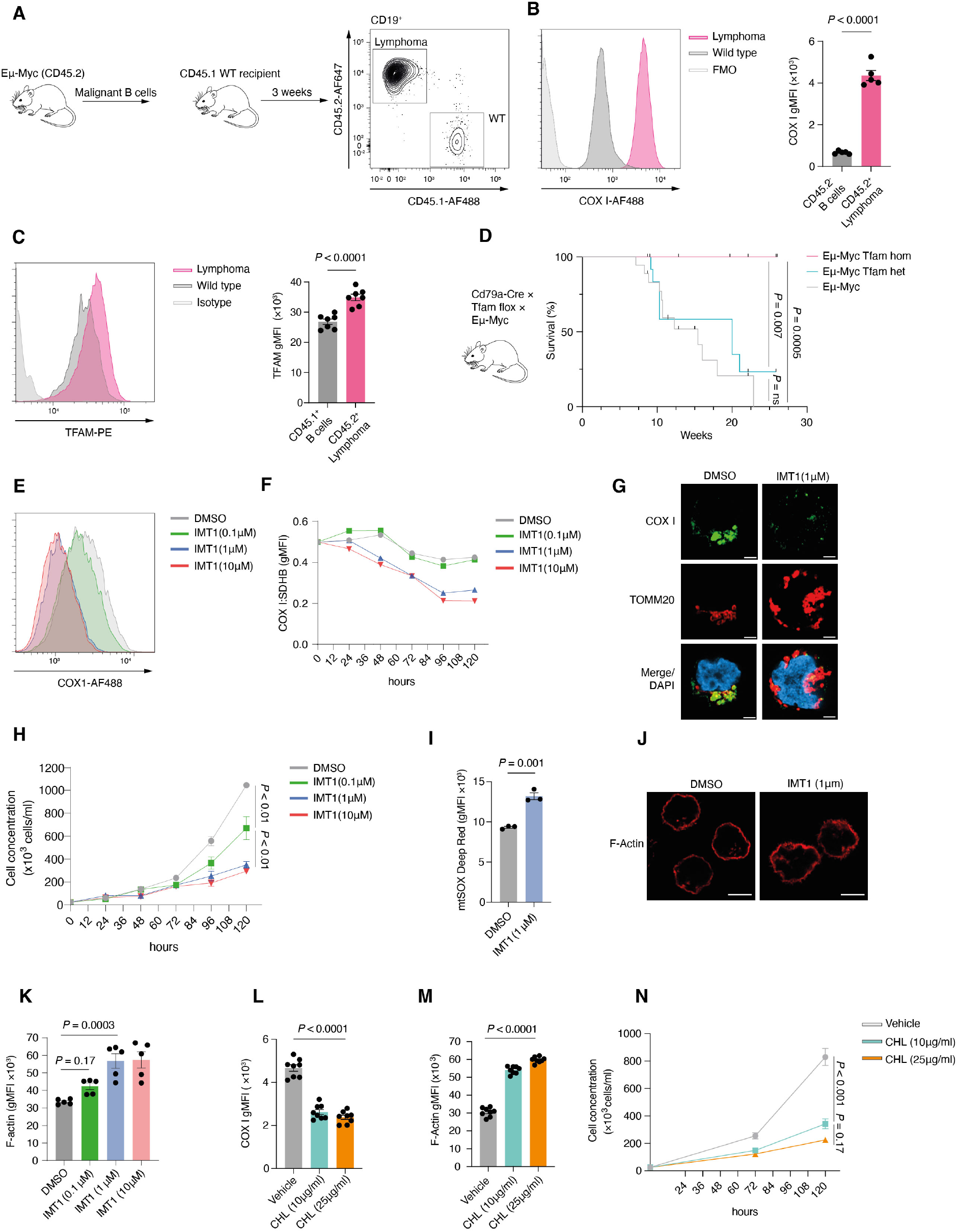
*Tfam* deletion in B cells abrogates the development and progression of lymphoma. A. Schematic depicting adoptive transfer strategy of lymphoma cells from Eμ-Myc mice. Following development of lymphoma, cells were transferred into wild type congenic (CD45.1^+^) recipients. Representative flow cytometry plots showing identification of CD19^+^CD45.2^+^ donor lymphoma cells and CD19^+^CD45.1^+^ wild type B cells from the inguinal lymph nodes of recipient mice after 3 weeks. B. Flow cytometry histogram depicting COX I expression in CD45.2^+^ lymphoma cells and CD45.1^+^ wild type B cells (n=5) as described in A. Quantification of COX I gMFI. Data representative of two independent experiments. C. Flow cytometry histogram depicting TFAM expression in CD45.2^+^ lymphoma cells and CD45.1^+^ WT B cells (n=5) as described in A. Quantification of TFAM gMFI. Data representative of two independent experiments. D. Kaplan-Meier graph depicting survival curve of Eμ-Myc × Tfam Hom (Cd79a^cre/+^ Tfam^fl/fl^ × Eμ-Myc, n=9), Eμ-Myc Tfam Het (Cd79a^cre/+^ Tfam^fl/+^ × Eμ-Myc, n=13) and Eμ-Myc (Cd79a^+/+^ Tfam^fl/fl^ × Eμ-Myc or Cd79a^+/+^ Tfam^fl/+ ×^ Eμ-Myc, n=18) mice followed for up to 6 months of age. E. Representative flow cytometry histogram depicting COX I expression in Daudi cell line treated with IMT1 at various concentrations (0.1μM, 1μM, 10μM) or vehicle (DMSO) for 120 hours. Representative of three independent experiments. F. Quantification of COX I levels by flow cytometry, normalised to SDHB in IMT1 or vehicle-treated Daudi cells at different time points (0, 24, 48, 72, 96, 120 hours), representative of two independent experiments. G. Airyscan ICC confocal images of COX I and TOMM20 in Daudi cells treated with DMSO or IMT1 (1μM) for 120 hours. Scale bar 3μm. Representative of two independent experiments. H. Quantification of live Daudi cell concentrations over 120h incubation period in the presence of IMT1 or vehicle, representative of two independent experiments with three technical replicates each. I. Flow cytometry-based quantification of mitochondrial ROS in Daudi cell line treated with IMT1 (1μM) or vehicle (DMSO) for 120 hours. Each point represents technical replicates. Representative of two independent experiments. J. Airyscan confocal images of Daudi cells treated with IMT1 (1μM) or DMSO, and stained for F-actin phalloidin. Scale bar 3μm. Representative of two independent experiments. K. Quantification of F-actin-phalloidin fluorescence by flow cytometry. Each point represents one technical replicate, pooled from two independent experiments. L. Quantification of COX I levels in Daudi cells by flow cytometry following treatment with vehicle (ethanol) or chloramphenicol at 10μg/ml or 25μg/ml for 120 hours. Data pooled from two independent experiments. M. Quantification of F-actin levels in Daudi cells by flow cytometry following treatment with vehicle (ethanol) or chloramphenicol at 10μg/ml or 25μg/ml for 120 hours. Data pooled from two independent experiments. N. Quantification of Daudi cell concentrations over 120-hour incubation period in the presence of chloramphenicol at 10μg/ml or 25μg/ml or vehicle (ethanol) as in M.Representative of two independent experiments with four technical replicates. Statistical significance was calculated by unpaired two-tailed t-test (B,C,I), Mantel-Cox log-rank test (D), two-way ANOVA with Šidák’s multiple comparison test (F), or ordinary one-way ANOVA with Dunnett’s multiple comparison test (H, K-N)

We next generated *Cd79a*-Cre × *Tfam* flox × Eμ-Myc mice (B-Tfam-Myc), to delete *Tfam* in B cells. B-Tfam-Myc mice were completely protected from the development of lymphoma during the observation period of 30 weeks, compared with *Tfam* flox × Eμ-Myc controls, which had a median survival of 10 weeks, or *Cd79a*-Cre × *Tfam* het × Eμ-Myc heterozygous mice (Fig. 8D).

Finally, to establish whether inhibition of mitochondrial transcription and translation might be a therapeutic target in human lymphoma, we treated Daudi B cell lymphoma cells (originally arising from Burkitt lymphoma) with IMT1, a specific inhibitor of mitochondrial RNA polymerase (POLRMT), which functions along with TFAM to initiate mitochondrial RNA transcription^55^. We found that treatment with IMT1 led to a progressive reduction of COX I levels, with increasing COX I:SDHB mismatch (Fig. 8E-F). Imaging of Daudi cells treated with IMT1 revealed mitochondrial enlargement, and confirmed the loss of COX I (Fig. 8G). IMT1 reduced cell proliferation, inhibited cell cycle progression, increased mitochondrial ROS levels in a dose dependent manner, and recapitulated the F-actin dysregulation we observed with *Tfam* deletion (Fig. 8H-K and Extended Data Fig. 8A). To inhibit mitochondrial translation we used chloramphenicol. Chloramphenicol reduced COX I expression, and increased expression of the UPR^mt^ protein LON peptidase 1 (LONP1), and F-actin in Daudi cells (Fig. 8L-M and Extended Data Fig. 8B). Cell growth was substantially reduced (Fig. 8N).

These results define high rates of mitochondrial translation enabled by *Tfam* expression as an essential requirement for the development of B cell lymphoma, and show the therapeutic potential of mitochondrial RNA polymerase and translation inhibition in human disease.

## Discussion

Here we show that upon entry to the GC, B cells dramatically remodel their mitochondria, increasing their mass and radically altering their morphology. As part of this transition, mitochondrial translation is highly active, and we demonstrate that the nuclear-encoded mitochondrial transcriptional and translational regulator TFAM is not only required for B cell development, but also for their entry into the GC program, proper spatial anchoring, and for the subsequent development of lymphoma.

GC B cells have a highly distinct metabolism, predominantly relying on OxPhos despite their very rapid rate of proliferation in a hypoxic microenvironment^6,8^. In order for rapidly dividing cells to maintain high mitochondrial mass in the face of dilution to their daughters, a high rate of mitochondrial protein translation and division must be maintained. We were surprised to find that TFAM is required for entry into the GC program itself, and when deleted there is a proportional accumulation of B cells with an activated precursor phenotype, which have expressed AID, but maintain markers of immaturity. It has recently been shown that the TCA metabolite ɑ-ketoglutarate is upregulated by IL-4 in B cells, and that this leads to epigenetic alteration of the *Bcl6* locus^9^. It is therefore possible that TFAM is required for an initial burst of mitochondrial biogenesis to facilitate GC program entry. Interestingly, GC B cells themselves have relatively reduced expression of TFAM compared to AP B cells, despite high levels of mitochondrial mass and dilution.

TFAM is responsible for packaging mitochondrial DNA into structures known as nucleoids, and also for maintenance of mtDNA copy number. The relative ratio of TFAM to mtDNA in nucleoids can dictate the expression of mitochondrial genes, based on the degree of genomic compaction^26^. It has been demonstrated that following activation in vitro, B cells have reduced mtDNA copy number and the number of nucleoids per mitochondrion drops, but their area increases^56^. In our transcriptional data, APs display generalised disruption of mitochondrial gene expression. In GC B cells however, mitochondrial transcription is generally reduced in the absence of *Tfam*.

Deletion of TFAM significantly altered the balance of ETC protein expression, with loss of mitochondrially-encoded subunits and compensatory upregulation of nuclear-encoded proteins. This was reflected in metabolic disturbance, with impaired OxPhos following activation, upregulation of glycolysis, and importantly, mitochondrial ROS generation, which was seen even in unstimulated cells. Consistent across genetic or pharmacologic interference with mitochondrial transcription and translation was an increase in F-actin. Dynamic actin cytoskeletal modification is essential for normal cell movement, and its probable disruption through ROS accumulation, reversible with the addition of a ROS scavenger, is likely to contribute to the positioning and motility defects we observed^57^. This adds weight to the idea that mitochondria are intimately linked to cytoskeletal function, and that this role may operate independent of ATP generation^58^. *Tfam* deletion also led to significant defects in calcium signalling, with upregulation of the MCU suggesting mitochondrial calcium sequestration. While not readily reversible with MCU inhibition, this may nonetheless also impair B cell function. Interestingly, B cells are especially sensitive to MCU deletion, illustrating their particular dependence on mitochondrial homeostasis^59^.

These defects collectively have the potential to compromise cellular interaction, in particular that with T_FH_ cells required to enter the GC program, and for normal dynamics once with in. We observed that Tfam^−/−^ B cells failed to differentiate normally into iGCs in an ex vivo T_FH_ co-culture system, and yet were unaffected when CD40 ligation was artificially provided. The capacity of Tfam^−/−^ iGB cells to present antigen to cognate T cells was normal however, as were T_FH_ cell numbers within GCs following immunisation. How *Tfam* deletion in B cells may affect T-cell interaction is therefore deserving of future study.

We did not observe an increase in GC B cell apoptosis with Tfam deletion, although it should be noted that due to highly effective efferocytosis by tingible body macrophages in the GC, differences may be challenging to detect.

Although we have directly established the importance of TFAM as a regulator of B cell development and activation, the factors driving the counterintuitive switch to OxPhos in GC B cells remain to be uncovered, as do the signalling mechanisms controlling the differences in mitochondria we observed between GC microenvironments. Disruption of mitochondrial integrity also induces a phenotype associated with immune aging, and to what extent the mechanisms we describe might hold true in the diminished humoral immune response seen with age is another area deserving of further exploration^20^.

We show that deletion of *Tfam* is sufficient to completely prevent the development of Myc-driven lymphoma. Although loss of Tfam at an early developmental stage leads to B cell lymphopenia, and therefore the pool of B cells which may become malignant is reduced, the high penetrance of the model contrasted with the complete protection against lymphoma suggests this is insufficient to explain the phenotype we observe. How TFAM acts to support lymphomagenesis requires further study, but may be associated with its promotion of mitochondrial translation^60^. Our observation that inhibition of mitochondrial transcription and translation reduces growth of lymphoma cells suggest that this should be prioritised as a therapeutic target.

## Methods

### Mice

B6.Cg-*Tfam^tm1.1Ncdl^*/J (JAX:026123), B6.C(Cg)-*Cd79a^tm1(cre)Reth^*/EhobJ (JAX: 020505), B6.129P2-*Aicda^tm1(cre)Mnz^*/J (JAX:007770), B6;129S6-*Gt(ROSA)26Sor^tm9(CAG-tdTomato)Hze^*/J (Ai9, JAX:007905), B6.Cg-Tg(TcraTcrb)425Cbn/J (JAX: 004194) and B6.Cg-Tg(IghMyc)22Bri/J [Eμ-Myc] (JAX: 002728) were purchased from Jackson Laboratories. Tg(Prdm1-Venus)1^Sait^ [Blimp1-mVenus] (MGI:3805969) was a kind gift from Mitinori Saitou (Kyoto University). Gt(ROSA)26Sor^tm1(CAG-mCherry/GFP)Ganl^ (MitoQC) was a kind gift from Ian Ganley (University of Dundee). B6.SJL.CD45.1 mice were provided by central breeding facility of the University of Oxford. Male and female mice between the ages of 6-15 weeks were used. Mice were bred and maintained under specific pathogen-free conditions at the Kennedy Institute of Rheumatology, University of Oxford. All procedures and experiments were performed in accordance with the UK Scientific Procedures Act (1986) under a project license authorized by the UK Home Office (PPL number: PP1971784).

### Immunisation

For SRBC immunisation, 1ml of sterile SRBCs (Fisher Scientific, cat: 12977755) were washed twice with 10ml of ice cold PBS and reconstituted in 3ml of PBS and administered as 0.2 ml injections intraperitoneally. In some experiments, an enhanced SRBC immunisation method was followed to maximise GC B cell yield by immunising mice with 0.1 ml SRBC on day 0 followed by 0.2 ml second injection on day 5^1^. For protein antigen immunisations, 50μg NP_(30-39)_-CGG (Biosearch Tech, cat: N-5055D-5) was mixed with Alum (ThermoFisher) or PBS for boost immunisation, at a 1:1 ratio and rotated at room temperature for 30 mins before intraperitoneal injection. For NP-CGG and SRBC-based immunisations, day 14 and day 12 were used as read-out time points respectively, unless specified otherwise.

### Flow cytometry and imaging flow cytometry

Flow cytometry was performed as previously described^2^. Briefly, harvested spleens were injected with ice cold PBS and mashed through a 70μm strainer (Falcon) or crushed between the frosted ends of microscope slides. For Peyer’s patch dissociation, a 40μm strainer (VWR) was used. RBCs were depleted by incubating splenocytes with ACK Lysis Buffer (Gibco) for 3-4 mins at RT. Single cell suspensions were incubated with Fixable Viability Dye eFluor™ 780 (eBioscience) in PBS, followed by FcBlock (5 mins) and surface antibodies (30 mins on ice) in FACS buffer (PBS supplemented with 0.5% BSA and 2mM EDTA). For intracellular staining, cells were fixed at RT with 4% freshly prepared PFA (Cell Signalling) for 15 mins and permeabilised with methanol (90% ice cold for 10 mins on ice with frequent vortexing) unless specified otherwise. Phalloidin-based F-Actin (Thermofisher, cat no: A30107) staining was performed using BD Perm/Wash reagent (BD, Cat. No. 554723) following 4% PFA fixation. For in vivo cell cycle analysis, 5-ethynyl-2’-deoxyuridine (EdU) (1 mg, Thermofisher, cat no: A10044) was injected intraperitoneally and mice were sacrificed after 2.5 h. Cells were stained for surface markers, fixed and permeabilised then labelled using Click chemistry according to manufacturer’s instructions (Click iT Plus EdU Flow cytometry kit, Thermo Fisher, cat: C10634). FxCycle Violet (Thermofisher, cat: F10347) reagent was used for cell cycle characterisation. For mitochondrial superoxide deep red (mtSOX, Dojindo) uptake, following viability dye and surface staining, cells were resuspended in warm complete RPMI supplemented with mtSOX (10μM) and incubated for 30 mins at 37°C. Cell were washed twice before flow cytometry acquisition. Flow cytometry was performed on BD Fortessa X-20 or LSR II instruments (both BD), or using a Cytek Aurora (5 laser) spectral flow cytometer. For imaging cytometry, single cell suspensions were prepared from spleens of MitoQC mice, incubated with live-dead dye, stained for surface markers, and then fixed with 4% PFA. Washed cells were then resuspended in 50μl FACS Buffer and run on an Amnis ImageStream^X^ Mark II Imaging Flow Cytometer and analysed with IDEAS (EMD Millipore) and FCS Express software (v7.12). Flow cytometry data was analysed using FlowJo (BD).

### Cell sorting

Naïve B cells were isolated using the Pan B cell Isolation II Kit, anti-CD43, and/or anti-CD23 Microbeads (all Miltenyi) according to the manufacturer’s instructions. Purity validated by flow cytometry was >90%. Isolation of DZ-LZ-GZ subsets of GC B cells, and T_FH_ cells (CD19^−^ CD4^+^ CXCR5^+^ ICOS^+^ GITR^−^) was performed via fluorescence-activated cell sorting (FACS).

For some experiments, untouched GC B cells were isolated using a magnetic bead-based protocol as described.^3^. Briefly, spleens were harvested from SRBC-immunised mice, and single cell suspensions were prepared in ice cold MACS isolation buffer (PBS with 0.5% BSA + 2mM EDTA) followed by ACK RBC lysis (Gibco) for 4 mins at RT with occasional mixing every 30s. After washing, cells were labelled with anti-CD43 microbeads (Miltenyi) and biotinylated antibodies against CD38 and CD11c (both eBioscience, clones 90 and N418 respectively), followed by incubation with anti-biotin microbeads, and subsequently run through a MACS LS column (Miltenyi). Purity was confirmed by flow cytometry and immunocytochemistry (ICC) and exceeded 95%.

When sorting for single cell RNA sequencing, spleens were crushed using the rough ends of microscope slides to maximise cell yield. Subsequently, non-B cells were depleted using the Pan B cell Kit II (Miltenyi). Enriched cells were then incubated with viability dye, anti-CD16/32 (FcBlock) and surface flow antibodies, including markers for an exclusion (dump) channel (anti-CD3, anti-Gr1 and anti-CD11c), and live Dump^−^ tdTomato^+^ cells were sorted by BD Aria FACS.

### Detection of ETC/UPR^mt^ proteins with spectral flow cytometry

A complete list of antibodies is shown in the antibody table. All antibodies targeting intracellular mitochondrial proteins were directly conjugated and of mouse origin, with the exception of rabbit anti-Tfam monoclonal antibody (Abcam). Goat anti-mouse AF405 Plus secondary antibody (ThermoFisher) was used for detection. All antibodies used in the panel had been either validated for flow cytometry by the vendor or in the literature. Mouse anti-mitochondrial complex 1 antibody was conjugated in house using a PE-Cy7 Lightning-link kit (Novus Bio, cat: 762-0005). Cells were labelled with Zombie NIR viability dye (Biolegend) and Fcblock in PBS for 30 min on ice in 96-well V-bottom plates. Following washing, surface staining was performed in Brilliant staining buffer (BD, cat: 563794) for 30 min. Cells were then fixed in 4% freshly-made PFA at RT for 15 mins and permeabilised in freezer-cold methanol for 10 mins with occasional vortexing. Anti-Tfam primary staining was performed in 50μl FACS Buffer supplemented with 2% goat serum for 45 mins at RT. Following two washes, goat anti-rabbit secondary antibody and the remaining directly conjugated antibodies for ETC/UPR^mt^ proteins were added for 30 mins at RT. The cells were subsequently washed in FACS buffer and acquired on a Cytek Aurora (5 laser) spectral flow cytometer. Exploratory pilot experiments were performed to determine the most suitable single-stained references (beads or cells) for individual marker-fluorochrome combinations. If cells were used as reference controls, they were obtained from matching organs (spleen or bone marrow). Reference controls were processed similarly to fully stained samples, with parallel fixation, permeabilization, and washing steps. Acquired samples were unmixed using SpectroFlow and analyzed with FlowJo. The ‘Autofluorescence (AF) as a fluorescent tag’ option was enabled during unmixing to minimise AF interference. In some rare cases minor adjustments were applied to unmixing on the SpectroFlow software. gMFI values for ETC/UPR^mt^ proteins were calculated using FlowJo.

### Flow cytometry antibodies

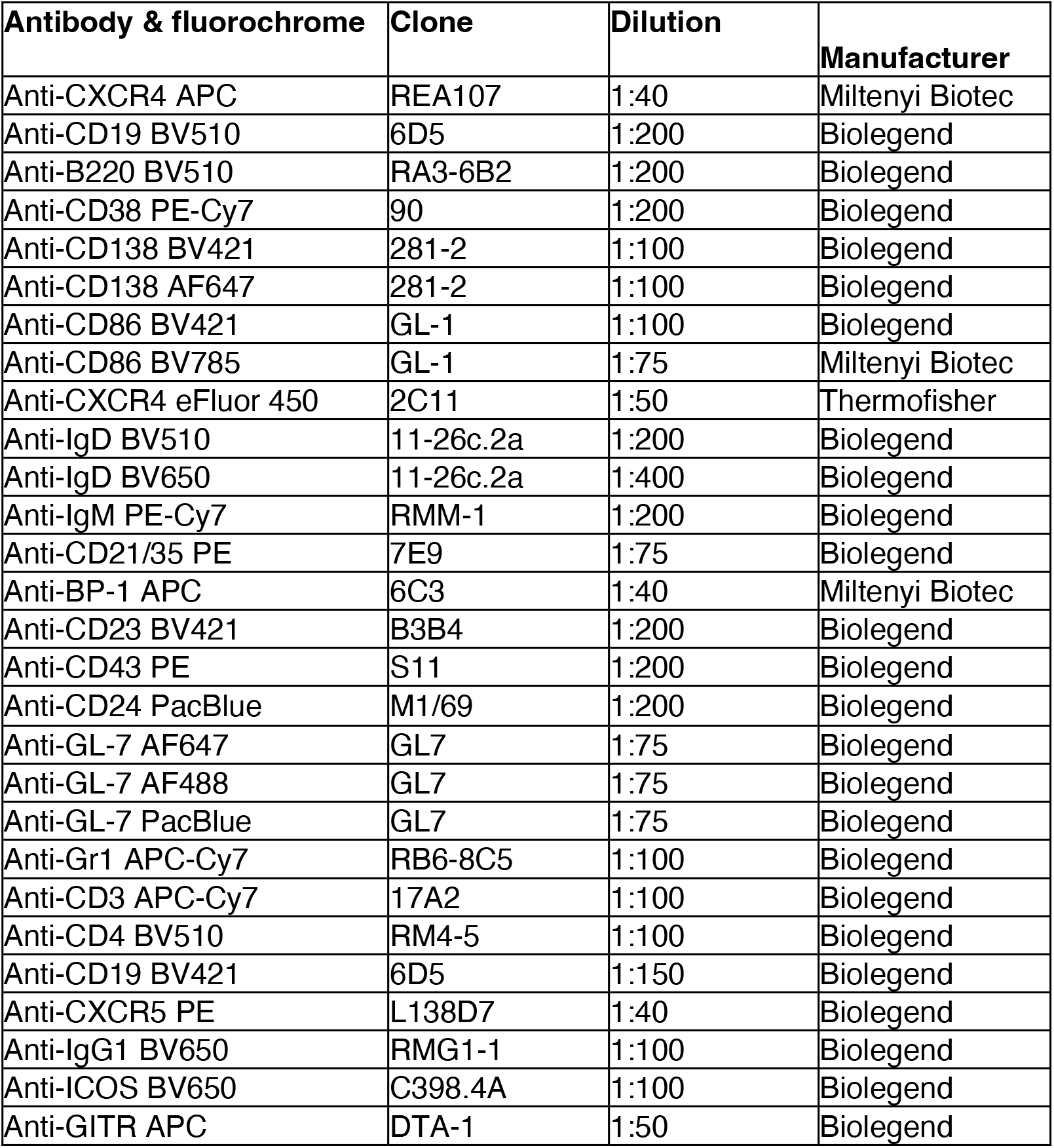
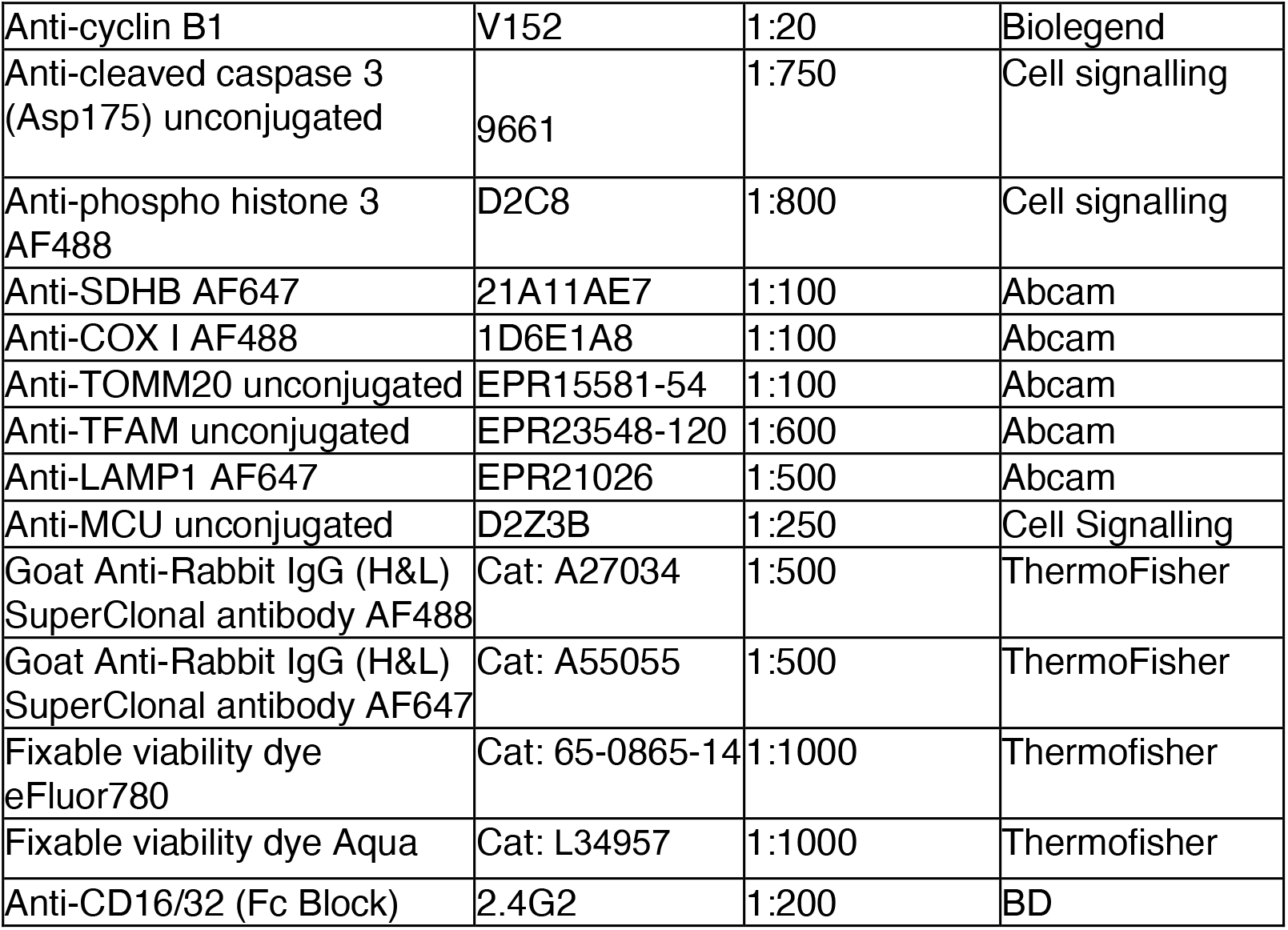

### Spectral flow cytometry antibody panel

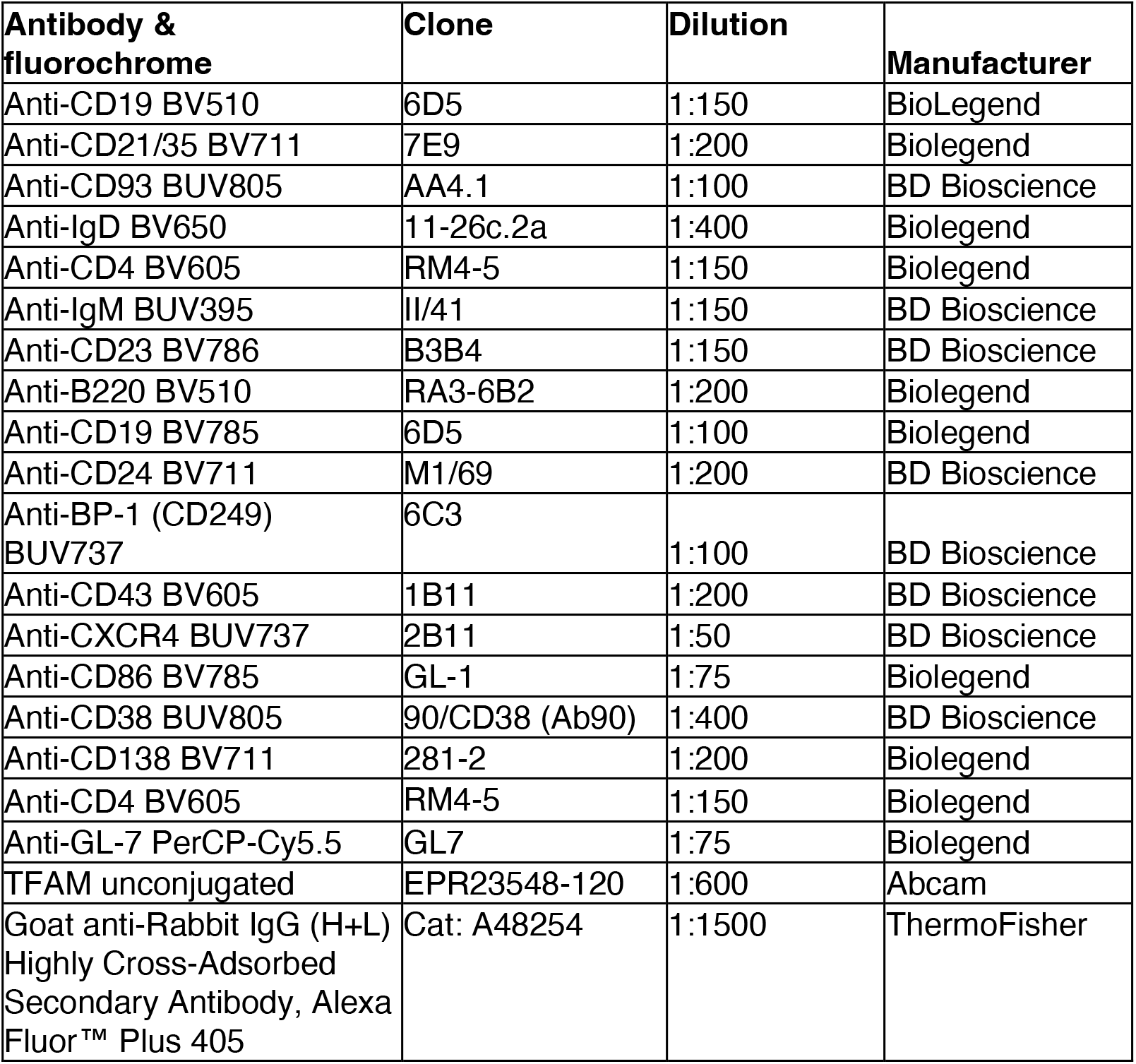
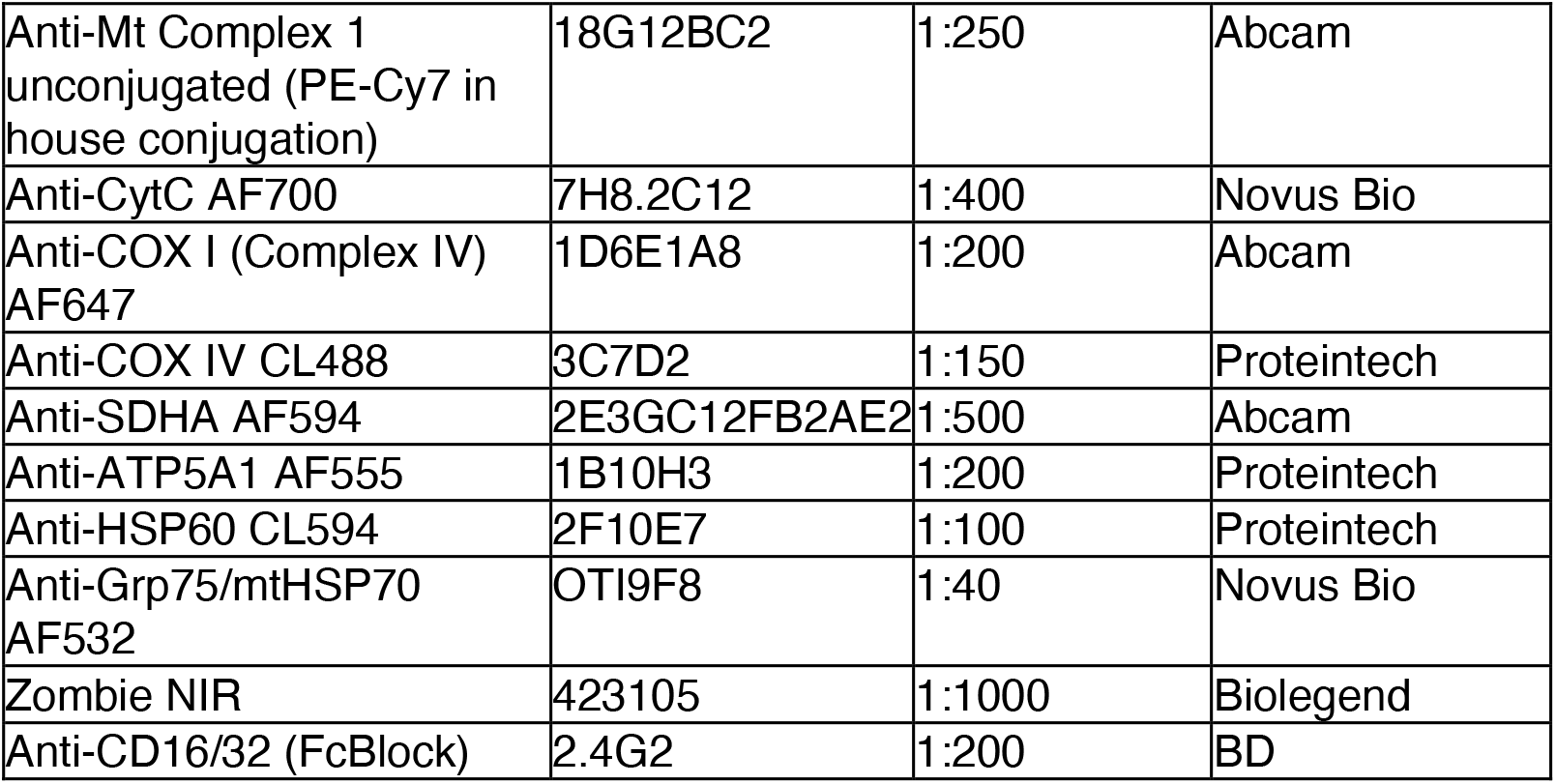

### Single-mitochondrion translation assay

The technique was performed as previously described, with modification^4^. Total B cells were isolated from spleens of Aicda-WT and Aicda-Tfam mice immunised with NP-CGG, using magnetic beads (Pan B cell Isolation Kit II, Miltenyi). Experiments were performed in batches such that one Aicda-WT and one Aicda-Tfam mouse spleen were processed in each replicate. Cells were pulsed with OPP (20μM, Jena Bioscience, cat: NU-931-5) in complete RPMI supplemented with the cytoplasmic translation inhibitor harringtonine (1μM, Abcam, cat: ab141941-5mg) for 45 mins at 37°C. Following a wash with complete medium, cells were resuspended in ice-cold Mitochondria Isolation Buffer (320mM sucrose, 2mM EGTA,10mM Tris-HCl, at pH 7.2 in water, prepared as 2× stock and stored in aliquots at −20°C). Cells were then homogenized with a Dounce homogenizer with a 2ml reservoir capacity (Abcam, cat: ab110169), using 10 strokes with type B pretzel. The homogeniser was rinsed with distilled water before each sample was processed to avoid cross-contamination. Homogenates underwent differential centrifugation at 4°C, with intact cells and isolated nuclei first pelleted at 1,000×g for 8 mins. The supernatant containing mitochondria was then transferred into new microcentrifuge tubes, and centrifugated at 17,000×g for 15 mins. Enriched mitochondria, which appeared as brown coloured pellets, were fixed in 1% PFA in 0.5ml PBS on ice for 15 mins, followed by a wash with PBS. Permeabilization and subsequent antibody staining was performed in 1× BD Perm/Wash buffer (diluted in MiliQ-filtered water) at RT. Pre-adsorbed primary rabbit anti-RFP (1:250, Rockland, cat: 600-401-379) antibody labelling was performed at RT for 30 mins. After a single wash, the Click reaction was performed using the Click-iT™ Plus OPP Alexa Fluor 647 Protein Synthesis Assay Kit (ThermoFisher, C10458) at RT for 30 mins. Following a wash, mitochondria were resuspended in 1× BD Perm/Wash buffer containing AF488 conjugated anti-COX IV (1:100, Proteintech, cat: 66110-1-Ig) and PE-conjugated Donkey anti-rabbit IgG (1:200, Biolegend, cat: 406421). Following wash with Perm/Wash buffer, mitochondria were resuspended in 250μl filtered (0.2μm) PBS and acquired using a BD Fortessa X20 flow cytometer. The threshold for SSC-A (log scale) was set to the minimum value (20,000) to allow acquisition of subcellular particles. Submicron Particle Size Reference Beads (Thermofisher, cat: F13839) were also used to identify mitochondria. For analysis, mitochondria were identified based on COX IV and SSC-A properties, and RFP^+^ (mitochondria with AP-GC origin) and RFP^−^ (mitochondria from naïve B cells) were gated based on anti-RFP signal. The gMFI for the OPP-AF647 fluorescence was calculated for RFP^+^ and RFP^−^ mitochondrial subsets, and their ratio was used as a translation index and pooled after batch correction. Mitochondria from reporter-free cells, and those untreated with OPP served as negative controls. In validation experiments 300μg/ml of chloramphenicol was used to inhibit mitochondrial translation along with the cytoplasmic translation inhibitor harringtonine, and mitochondria were detected by COX IV-AF488 and SDHA-AF594 antibodies (1:250, Abcam, ab170172) as described above.

### NP conjugation

NP-APC conjugation was performed as described^5^, with certain modifications. Briefly, one mg of natural allophycocyanin protein (APC) (Stratech Scientific Ltd) was transferred into Slide-A-Lyzer™ MINI Dialysis Device, 3.5K MWCO (Thermofisher, cat: 69550) and dialysed for 5 hours, overnight, then for 4 hours in 1L 3% NaHCO_3_ at 4°C. NP-Osu (Biosearch, cat no: N-1010-100) was dissolved in dimethylformamide (DMF) (Merck) to a concentration of 10mg/ml while vortexing. The NP-Osu was added to the dialysed APC at a ratio of 20μg:1 mg (NP-low conjugation) and 80μg:1 mg (NP-high conjugation) and rotated at RT for 2 hours, protected from light. The NP-APC conjugates were then dialysed in 1L 3% NaHCO_3_ at 4°C overnight, then 1L PBS overnight. NP probes were stored at 4°C in the dark until use.

### ELISA

96 well EIA/RIA plates (Corning, cat: 3590) were coated with NP-BSA (NP_1-4_ or NP_<20_, cat: N-5050XL-10 and cat: N-5050H-10) at 5μg/ml in bicarbonate/carbonate coating buffer overnight at 4°C. The next day, plates were washed with PBS and blocked with 5% skimmed milk in PBS for 2.5 hours at 37°C. Sera obtained from mice (NP-CGG-immunised day 14 & 49) were serially diluted in 1% skimmed milk and incubated in blocked plates at serial dilutions for 1 hour at 37°C. After multiple washes with PBS supplemented with 0.05% Tween-20, alkaline phosphatase-conjugated goat anti-mouse IgG1 or IgM (Southern Biotech, cat: 1071-04, cat: 1021-04) detection antibody was added (1:2000) and incubated for 1 hour at 37°C. After the final washing step, plates were developed with AP substrate (Sigma, cat: P7998-100ML) for 10-30 mins and read on a FLUOstar Omega plate reader at 5 min intervals.

### IHC

Harvested spleens were immediately transferred to Antigenfix (DiaPath) solution and fixed overnight at 4°C. The next day, spleens were washed in PBS followed by overnight incubation in 30% sucrose (in PBS) at 4°C for cryoprotection. On the following day, spleens were snap frozen in 100% methanol on dry ice and stored at −80°C until cryosectioning at 8-12μm thickness. Slides were then rehydrated in PBS at RT, then permeabilised and blocked in PBS containing 0.1% Tween-20, 10% goat serum, and 10% rat serum at RT for two hours. For panels requiring intracellular detection, Tween-20 was replaced by 0.5% Triton X-100. All primary antibody staining was performed overnight at 4°C in PBS supplemented with 2% goat serum and 0.1% Triton X-100 (intracellular) / Tween-20 (surface), and secondary staining was performed at RT for 2 hours the next day in PBS 0.05% Tween-20. TUNEL imaging was performed using the Click-iT Plus TUNEL In Situ Imaging Far Red kit according to manufacturer’s protocol (Thermofisher, cat: C10619). Slides were mounted with Fluoromount G (Southern Biotech, cat: 0100-01) and imaged using a Zeiss LSM 980 equipped with an Airyscan 2 module.

### ICC

Isolated cells were transferred onto 18mm coverslips coated with 0.001% poly-L-lysine (Merck, cat: P4707-50ML, 10× stock) and incubated for 10 mins at 37°C to ensure cell attachment. Cells were then fixed in prewarmed 1× PHEM buffer (60mM PIPES, 25mM HEPES, 10mM EGTA, and 4mM MgSO4·7H20) with 4% PFA for 10 mins at 37°C, followed by permeabilization and blocking in 0.2% Triton X-100 with 10% goat serum for 30 mins. Primary antibody labelling was performed overnight at 4°C, and secondary antibody staining for 45 mins at RT (see antibody table). For the mitochondrial transcription assay based on 5-ethynyl uridine (EU) incorporation, isolated untouched naïve B cells and GC B cells were resuspended in complete RPMI supplemented with 1mM 5-EU and transferred to 18mm coverslips coated with poly-L-lysine. Following incubation for 45 mins, the cells were briefly washed and fixed in warm 4% PFA diluted in PHEM Buffer. Following permeabilization and blocking for 30 min, incorporated 5-EU was detected by the Click-iT™ RNA Alexa Fluor 594 Imaging Kit (ThermoFisher, cat: C10330). Intracellular antibody labelling was performed after the Click reaction, to minimise the interference of Click reagents with fluorochromes. For in vivo measurement of RNA synthesis, 2mg of 5-ethynyl uridine (5-EU, Thermofisher) was injected intraperitoneally at D12 following SRBC immunisation and similar preparation and labelling steps described for the ex vivo 5-EU assay were followed. 4′,6-diamidino-2-phenylindole (DAPI, Sigma, cat: D8417-1MG) staining was performed at 1μM at RT for 5 mins, followed by a brief wash and mounting in Fluoromount G (Southern Biotech). For Mitotracker staining, cells were labelled with Mitotracker Red CMX ROS (150nM, Thermofisher, cat: M7512). Slides were mounted with Fluoromount G (Southern Biotech, cat: 0100-01) and imaged using a Zeiss LSM 980 equipped with an Airyscan 2 module.

### STED microscopy

STED super resolution imaging was performed using a Leica TCS SP8 laser scanning STED system equipped with an oil objective (HC PL APO 100× NA 1.40) and a 775nm depletion laser. Isolated naïve and GC B cells labelled with anti-TFAM (primary, Abcam) and Alexa Fluor 647 conjugated goat anti-rabbit secondary antibody (ThermoFisher, cat: A55055) were imaged in confocal and STED mode sequentially. COX I labelled with AF488 was subsequently imaged in confocal mode to define mitochondrial boundaries. Acquired STED images were deconvoluted using Deconvolution Express mode with Standard setting in Huygens Essential software (v22.04) (Scientific Volume Imaging, Hilversum, Netherlands) and exported using Image J software.

### IHC & ICC

| Antibody & fluorochrome | Clone | Dilution | Manufacturer |
| --- | --- | --- | --- |
| Anti-CD38 AF647 | 90 | 1:50 | Biolegend |
| Anti-CD21/35 AF647 | 7E9 | 1:75 | Biolegend |
| Anti-CD21/35 FITC | 7E9 | 1:75 | Biolegend |
| Anti-CD138 AF647 | 281-2 | 1:50 | Biolegend |
| Anti-IgD AF647 | 11-26c.2a | 1:50 | Biolegend |
| Anti-GL-7 AF647 | GL-7 | 1:50 | Biolegend |
| Anti-GL-7 AF488 | GL-7 | 1:50 | Biolegend |
| Anti-GL-7 PacBlue | GL-7 | 1:50 | Biolegend |
| CD3e Monoclonal Antibody | 500A2 | 1:100 | eBioscience |
| Anti-SDHB AF647 | 21A11AE7 | 1:75 | Abcam |
| Anti-COX I AF488 | 1D6E1A8 | 1:100 (IHC)<br>1:500 (ICC) | Abcam |
| Alexa Fluor 647 Goat Anti-Hamster IgG (H+L)-0.5 mL | Cat: A-21451 | 1:1000 | Thermofisher |
| Goat Anti-Rabbit IgG (H&L) SuperClonal antibody AF488 | Cat: A27034 | 1:500 | ThermoFisher |
| Goat Anti-Rabbit IgG (H&L) SuperClonal antibody AF647 | Cat: A55055 | 1:500 | ThermoFisher |
| Anti-TOMM20 unconjugated | EPR15581-54 | 1:500 | Abcam |
| Anti-Lonp1 unconjugated | HPA002192 | 1:300 | Atlas |
| Anti-TFAM unconjugated | EPR23548-120 | 1:500 | Abcam |

### Image analysis

Zen Blue (v3.4, ZEISS) and ImageJ software were used for image analysis/processing. For the determination of GC properties in splenic sections, tdTomato and/or GL-7 signals were used as a reference to identify GCs, and IgD for naïve B cell follicles. Defined areas were introduced as regions of interest (ROIs) using the Analyse Particles function. Mitochondria were segmented based on MitoQC GFP or COX I signal, and subsequently area and signal intensity calculations were performed using the Analyse Particles function in ImageJ. For 3D volumetric analyses of mitochondria, the 3D Objects Counter function was used. Mitochondrial 5-EU incorporation is quantified first by identifying mitochondrial area based on thresholded COX I signal. Background noise in the 5-EU channel was removed using the Subtract Background (Rolling ball radius 20 pixels) function. Areas outside of the mitochondrial ROIs were removed, and the remaining 5-EU integrated signal density was determined by the 3D Objects Counter function. 3D TFAM-mitochondrial nucleoid complexes were enumerated by 3D Suite^6^. Briefly, local maxima of TFAM signals were determined for each z-slice using 3D Fast Filters (kernel x, y and z 1px each) which were then inputed into a 3D Spot segmentation module with local thresholded Gaussian fit (Radius max 10 px, SD value 1.50).

PC-PB quantification on splenic IHC images was performed in ImageJ software. Confocal images were imported in split channel mode. For total splenic section area quantification, the DAPI channel was thresholded (using Yen setting) and measured via the Analyse Particles function (20μm size threshold + include holes). Then, a Gaussian blur filter (2 sigma radius) was applied to CD138, tdTomato and Blimp1-mVenus channels followed by Background subtraction (rolling ball radius 50px). Subsequently, each channel was autothresholded (Yen) and Watershed segmentation was applied to thresholded binary images. CD138 and tdTomato channels were inputted into Image calculator using the AND function. The resulting image was then inputted into the same mode, including Blimp1-mVenus channel. The final image was subsequently quantified using the Analyse Particle mode (0.5μm size threshold + Include holes) to identify the triple positive PC fraction. PB numbers were then calculated as follows: CD138^+^ Blimp1-mVenus^+^ cell number minus triple positive (CD138^+^ Blimp1-mVenus^+^ tdTomato^+^) PC number. For T_FH_ enumeration on splenic sections, first tdTomato^+^ GC clusters were auto-thresholded (Otsu). GC area was calculated using the Analyse Particle function. After brightness/contrast adjustments, CD3^+^ T_FH_ cells within GC ROIs were manually counted.

For DZ/LZ analyses in tissue sections, the nesting function in Zen Blue was used by identifying tdTomato^+^ GCs as ROIs. The CD21/35 signal was then used to calculate the area of the LZ. Normalised DZ area was quantified as follows: (GC area – LZ area) / GC area. Quantification was performed for each individual GC pooled from splenic sections. GCs with values >0.95 were excluded as not including any representative LZ area. Airyscan reconstruction was performed in Zen Blue software with Medium filter setting for images acquired with the relevant module. ImageJ Macro codes used for image analyses are available upon request.

When GC B cells were isolated from Aicda-Cre × Rosa26^STOP^tdTomato mice, the tdTomato reporter signal was used to filter out contaminating non-GC B cells.

### Quantification of metabolism by OPP incorporation

This technique was performed as described, with modification^7^. Briefly, splenocytes were split into 4 groups and incubated for 30 mins with or without metabolic inhibitors (1μM oligomycin and/or 1mM 2DG, both from Merck) and their combinations in 96 well plate at 37°C. The alkynylated puromycin analog OPP (20μM final concentration, ThermoFisher) was then directly added to the wells for an additional 30 mins incubation. Subsequently, cells were washed and labelled with Live/Dead viability dye and surface antibodies, after which they were fixed with 4% PFA. Click Chemistry labelling was performed according to Click-iT™ Plus OPP Alexa Fluor 647 Protein Synthesis Assay Kit (ThermoFisher, cat: C10458). The percentages of mitochondrial dependence, glycolytic capacity, glucose dependence and FAO/AAO (fatty acid oxidation and amino acid oxidation) were measured using the gMFI values of OPP-AF647 of cells treated with 2-deoxyglucose (2-DG), oligomycin, or 2-DG and oligomycin. The formulae are summarised below:

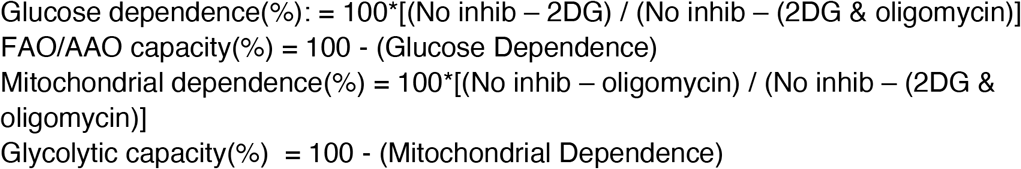

### Extracellular flux analysis

Real-time oxygen consumption rates (OCR) and extracellular acidification rates (ECAR) were recorded using a Seahorse XFe96 Extracellular Flux Analyzer (Agilent). Briefly, ex vivo isolated naïve or overnight stimulated (anti-CD40 at 5μg/ml and IL-4 at 1ng/ml) 3×10^5^ B cells were plated on a poly-D lysine (PDL) (Sigma)-coated XF96 cell culture microplate and incubated at 37 °C for a minimum of 30 min in a CO_2_-free incubator in assay medium (XF RPMI medium pH 7.40 supplemented with 2 mM L-glutamine, 1 mM pyruvate and 10 mM glucose). iGB cells were rested in the presence of IL-4 at 1ng/ml overnight after detachment from a fibroblast feeder layer. The next day, they were plated at 2 × 10^5^ per well density on PDL-coated XF96 cell culture microplate. Basal OCR and ECAR were measured, then followed by the addition of the MitoStress Test inhibitors: oligomycin (1μM), fluorocarbonyl cyanide phenylhydrazone (FCCP, 2μM) and rotenone + antimycin A (0.5μM) (Agilent). Analyses were performed on Agilent Wave and Prism software.

### In vitro mouse primary B cell culture

B cells were isolated from spleens of unimmunised Tfam-floxed and wild type mice carrying the Rosa26-stop-tdTomato allele without Aicda-Cre using the Pan B cell isolation kit II (Miltenyi). TFAM was deleted by TAT-Cre recombinase (1.5μM, Merck, cat: SRC508) in serum-free complete RPMI (supplemented with 2mM GlutaMAX, 1mM pyruvate, 10mM HEPES, 100 IU/ml Penicillin/Streptomycin and 50μM 2-mercaptoethanol) for 45 mins at 37°C and 5% CO_2_. CellTrace Violet dye (Thermofisher) at 10μM was added directly after 30 mins of TAT-Cre incubation and cells were incubated for an additional 15 min at 37°C. Subsequently, cells were washed three times with 10% FBS containing complete RPMI, and live cells were counted manually using Trypan Blue exclusion of dead cells. B cells were cultured in U-bottom 96-well plates at a concentration of 1×10^6^/ml with anti-IgM (1μg/ml, Jackson Immuno), anti-CD40 (1μg/ml, Miltenyi) and IL-4 (40ng/ml, Peprotech) stimulation for four days at 37°C in a humidified incubator with 5% CO_2_.

### iGB culture system

The iGB culture system was described previously by Nojima et al^8^. Briefly, the 3T3 fibroblast cell line of BALB/c origin stably expressing CD40 ligand (CD40L) and B-cell activating factor (BAFF) (40LB cell line), was cultured and maintained in high glucose DMEM with GlutaMAX (Life Technologies, cat: 31966021) medium supplemented with 10% FBS and 100IU/ml penicillin/streptomycin (Life Technologies, cat: 15070063) in T75 tissue culture flasks (Sarstedt, cat: 658175). Once cells were confluent, they were detached using trypsin/EDTA (Gibco, cat: 25200056) treatment, washed, and collected in 15 ml tubes in 5 ml medium and irradiated (80 Gy). Following irradiation, the cells were washed, counted and seeded at 3×10^6^ per dish (100mm, Greiner bio, G664160) and 0.5×10^6^ per well (6 well plate, Falcon, 353046), respectively. Fibroblast attachment and stretching were allowed overnight at 37°C and 5% CO_2_. The next day, naïve B cells were isolated using anti-CD43 micro-beads and treated with TAT-Cre (~1.5μM) for 45 mins as described above. Following three washes, the cells were counted and cultured on an irradiated 40LB layer at 5×10^5^ (100 mm dish) and 5×10^4^ (per well, 6 well plate) for 4-6 days.

### iGB adoptive transfer

iGB cells were generated as above. On day 4, fibroblasts and in vitro differentiated plasmablasts (generally <10% frequency) were removed using biotinylated anti-H-2Kd (Biolegend, cat: 116604) and anti-CD138 (Biolegend, cat: 142511) followed by anti-biotin microbeads (Miltenyi) and negative selection was performed using LS columns. In some experiments, FACS was used for the purification of tdTomato^+^ iGB cells. For competitive experiments, purified wild type iGBs (CD45.1/2^+^) were mixed 1:1 (ratio confirmed by flow cytometry prior to transfer) with Tfam^−/−^ iGBs and injected intravenously (6×10^6^ cells in competitive settings or 3×10^6^ cells in non-competitive settings) into CD45.1^+^ or CD45.2^+^ congenic hosts that were immunised with SRBC according to the enhanced protocol to maximise the recruitment of transferred iGB cells into the GC. On day 6 post transfer, spleens were harvested and analysed by flow cytometry and confocal imaging to assess GC entry.

### T_FH_-B cell co-culture

The method was described by Sage et al^9^. Briefly, CD19^−^CD4^+^CXCR5^hi^ICOS^+^GITR^−^ T_FH_ cells were isolated from SRBC-immunised (enhanced protocol) wild type mice at day 14. Naïve B cells were isolated from *Tfam flox* and wild type mice, both carrying the Rosa26^STOP^tdTomato allele with anti-CD43 microbeads (Miltenyi). Following TAT-Cre treatment (~1.5μM, 45 min), 5×10^4^ wildtype or Tfam^−/−^ B cells were co-cultured with 3×10^4^ T_FH_ cells in U bottom 96 well-plate in the presence of anti-CD3 (2μg/ml) (145-2C11, ThermoFisher, cat: 16-0031-82) and anti-IgM (5μg/ml, Jackson Immuno) for 6 days. At the end of the culture period, the percentages of CD19^+^ GL-7^+^ IgG1^+^ tdTomato^+^ iGCs were quantified via flow cytometry.

### Peptide presentation and T-B conjugate assay

The technique was performed as previously described^10,11^, with modifications. CD45.1^+^ GFP^+^ OT-II CD4^+^ T cells were purified from mouse spleens using magnetic beads (CD4 T cell MojoSort isolation kit, Biolegend) and activated with 10μg/ml recombinant mouse IL-2 IS (Miltenyi, 130-120-662) for 3 days in 24-well plates coated with 3μg/ml anti-CD3 (145-2c11; Biolegend) and 3μg/ml anti-CD28 (37.51, Biolegend). Naïve B cells were isolated from Rosa26^STOP^tdTomato × Tfam WT or Tfam flox mice using anti-CD43 micro-beads (Miltenyi), and following TAT-Cre administration, activated for 4 days using the iGB-40LB in vitro system in 6 well plates. On day 4, fibroblasts were removed, and 2×10^5^ purified iGB cells were pulsed with ovalbumin peptide (amino acids 323–339) (Genescript) at varying concentrations (0μM, 0.1μM and 1μM) for 30 min at 37°C in 96 well U-bottomed plates (Falcon) in triplicates per condition. After the addition of 5×10^5^ activated OT-II cells to each well, the plates were centrifuged at 500×g for 5 min at 25°C. The centrifuged cell pellets were then incubated for 30 min at 37°C. 16% PFA was added directly to the wells to achieve a final concentration of 4%, and the pellets were gently resuspended. The cells were fixed for 10-15 mins and directly acquired using an Aurora spectral flow analyser (Cytek) at low flow rate setting to minimise artificial doublets. 15,000 events were recorded per condition. Before acquisition, the plate was shaken at 1100rpm by the plate holder. As a negative control, T and B cells were mixed in the absence of OVA peptide without centrifugation and subsequent incubation.

### Transwell migration assay

5×10^5^ enriched total B wells isolated from SRBC-immunised Aicda-Tfam and Aicda-WT mice were placed in a 6mm transwell chamber with 5μm pore size (Corning, cat: 3421) and incubated at 37°C for two hours in the presence of murine CXCL12 (200ng/ml, Biolegend) or CXCL13 (1μg/ml, Biolegend) in complete RPMI. Relative migration index was calculated as follows: percentage of GC B cells (CD38^−^GL-7^+^tdTomato^+^) in total input cells divided by the percentage of GC B cells in migrated live total cells. 2×10^5^ purified total B wells from B-Tfam mice placed in a 6mm transwell chamber with 5μm pore size and incubated for five hours in the presence or absence of murine CXCL12 (200ng/ml, Biolegend) with or without mitoTempo (100μM, Merck) and Ru265 (30μM, Merck). After five hours, cells were incubated with Live/Dead and anti-B220 AF488 antibody, and resuspended in 100μl in 96 well V bottom plates and acquired on a Cytek Aurora flow cytometer at high flow setting with a stopping volume of 60μl. Technical duplicates were also included. Specific migration (%) was calculated following this formula: 100 * [No. of B220^+^ cells migrated in response to CXCL12 – No. of B220^+^ cells migrated in the absence of CXCL12]/No. of input B cells.

### Cytoplasmic calcium assay

Due to the time and temperature sensitive nature of calcium dyes, experiments were performed in four batches such that one wild type and one B-Tfam mouse spleen were processed in each replicate. Single cell suspensions were prepared from spleens and 2×10^6^ cells were placed in V bottom 96 well plates, then labelled with fixable viability dye and then anti-B220 antibody. Subsequently, cells were stained with the cytoplasmic calcium indicator dye Fluo4-AM (Thermofisher, cat: F14201) at 10μg/ml in Iscove’s Modified Dulbecco Media (IMDM) supplemented with 1% FBS at 37°C for 15 mins. Following a single wash, cells were resuspended in 400μl warm IMDM with 1% FBS. Samples were kept in a warm water container (~37°C) throughout the flow cytometry acquisition to maintain physiologic activity. Baseline fluorescence intensity was measured for 30s prior to stimulation with either CXCL12 (200ng/ml) or anti-IgM (10μg/ml). Flow cytometry acquisition was performed on a BD Fortessa X20 at low speed setting for 5 min. The acquisition sequence was alternated between wild type and experimental samples in each batch to avoid potential timing-related noise. Analyses were performed using the Kinetics module in Flow Jo software.

### Lymphoma adoptive transfer

Following the manifestation of clinical signs of lymphoma, Eμ-Myc mice were sacrificed and spleens were harvested. Non-B cells were depleted using the Pan B cell Isolation Kit II (Miltenyi) to enrich lymphoma cells. 2×10^6^ lymphoma cells were then intravenously injected into recipient B6.SJL.CD45.1 mice in 200μL PBS. Recipient mice were sacrificed at 3 weeks following adoptive transfer, and spleens and inguinal lymph nodes were harvested for further analyses.

### Mixed bone marrow chimera generation

B6.SJL.CD45.1 recipient mice were given two doses of 5.5Gy irradiation four hours apart. Mice were then intravenously injected with 4×10^6^ mixed bone marrow (BM) cells at a 1:1 ratio, isolated from age- and sex-matched CD45.2^+^ Aicda-WT and CD45.1^+^ WT or CD45.2^+^ Aicda-Tfam and CD45.1^+^ WT donor mice. Recipient mice were maintained on antibiotics (Baytril, Bayer corporation) in drinking water for two weeks. BM reconstitution was confirmed via flow cytometry of peripheral blood at 8 weeks. Mice were immunised with SRBC at 11 weeks following irradiation and analysed at day 7.

### Lymphoma cell culture system

Daudi cells were cultured in RPMI 1640 medium supplemented with 10% FBS, 1× GlutaMAX (Gibco), 25mM HEPES and 100U/ml penicillin and 50μg/ml streptomycin and maintained at 37°C in a humidified incubator with 5% CO_2_. IMT1 (MedChem Express, as a 1mM stock solution in DMSO, cat no. HY-134539) was used at 0.1μM, 1μM, and 10μM concentrations for a 0-120h time window. Chloramphenicol (Merck, cat no: C0378-5G) was used at 10μg/ml and 25μg/ml concentrations (prepared in 100% ethanol fresh for each culture experiment) for a 0-120h time window. Cell numbers were determined by manual counting using Trypan blue dye for dead cell exclusion at each timepoint.

### Single cell RNA-seq analysis

Approximately 17,000 cells per sample were loaded onto the 10x Genomics Chromium Controller (Chip K). Gene expression and BCR sequencing libraries were prepared using the 10x Genomics Single Cell 5’ Reagent Kits v2 (Dual Index) following manufacturer user guide (CG000330 Rev B). The final libraries were diluted to ~10nM for storage. The 10nM library was denatured and further diluted prior to loading on the NovaSeq6000 sequencing platform (Illumina, v1.5 chemistry, 28bp/98bp paired end) at the Oxford Genomics Centre.

Filtered output matrices from Cellranger 6.0.1 were loaded in *Seurat* v4.1.0. Cells with more than 5% mitochondrial reads and <200 genes were removed from analysis. Data were normalised and transformed using SCTransform, with regression of cell cycle phase and mitochondrial reads, and integrated using FindIntegrationAnchors and IntegrateData functions. Principal component analysis and UMAP were used to cluster cells. Marker genes between clusters were identified using the FindAllMarkers function. Two small contaminant clusters (<1% of cells) were identified based on expression of non-B cell genes and were removed from subsequent analysis. Clusters were identified by expression of canonical markers. Cluster proportions were calculated using *DittoSeq*. For visualisation of UMAP projections, equal number of cells from each experimental condition were displayed by random downsampling. Differential gene expression between conditions was calculated using the FindMarkers function with the ‘t-test’ parameter. Pathway analysis was performed using *SCPA*. Pseudobulk differential gene expression between individual biological replicates was performed using *EdgeR* after count aggregation across cells using *Scuttle*.

Filtered contig outputs generated by Cellranger 6.0.1 from cells processed in the Seurat workflow above were combined, filtered, and visualised using *scRepertoire* 1.5.2. For quantification of mutational load, the *Immcantation* pipeline was employed. Germline segment assignment was performed using *Change-O*, and SHM count was calculated using *SHazaM*.

### Statistical analysis

The use of the statistical tests is indicated in the respective figure legends, with the error bars indicating the mean ± S.E.M. P values ≤ 0.05 were considered to indicate significance. Analyses were performed with GraphPad Prism v9 or R 4.1. No formal power calculations were performed. The distribution of data was determined using normality testing to determine appropriate statistical methodology. In selected experiments batch correction was performed using the R package *Batchma*.

## Data and code availability

Single cell RNA sequencing data has been deposited to GEO under accession number GSE208021. Code used in analysis of data is available upon reasonable request.

## Acknowledgements and funding

We thank Jonathan Webber for flow sorting, and Tal Arnon and Peter McGill (University of Oxford) for providing mice and Lynn Dustin (University of Oxford) for providing Daudi cells. We thank Daisuke Kitamura (Tokyo University of Science) for providing the 40LB cell line. We also thank Karl Morten (University of Oxford) for helpful suggestions.

Funding for this work was provided by the Wellcome Trust (211072/Z/18/Z) and Cancer Research UK/Versus Arthritis (C70663/A29547) to A.J.C., the Kennedy Trust for Rheumatology Research, and the US National Institutes of Health (HL118979) to M.L.D. S.J.D. is funded by an NIHR Global Research Professorship (NIHR300791). Flow cytometry and microscopy facilities were supported by the Kennedy Trust for Rheumatology Research through the Cell Dynamics Platform. We thank the Wolfson Imaging Centre Oxford for providing microscope facility support, and the Don Mason flow cytometry facility of the Dunn School, University of Oxford.

## >Author contributions

Y.Y. and A.C. conceived and designed the study. Y.Y. performed the majority of the experiments. E.M., E.C., S.G., C.S., M.A., B K-D, and M.A. performed experiments. Y.Y. and A.C. wrote the manuscript. Y.Y. performed image analysis and A.C. and I.R. analysed single cell data. M.D. and S.D. provided advice and guidance. A.C. supervised the study.

---

**Extended Data Figure 1.**
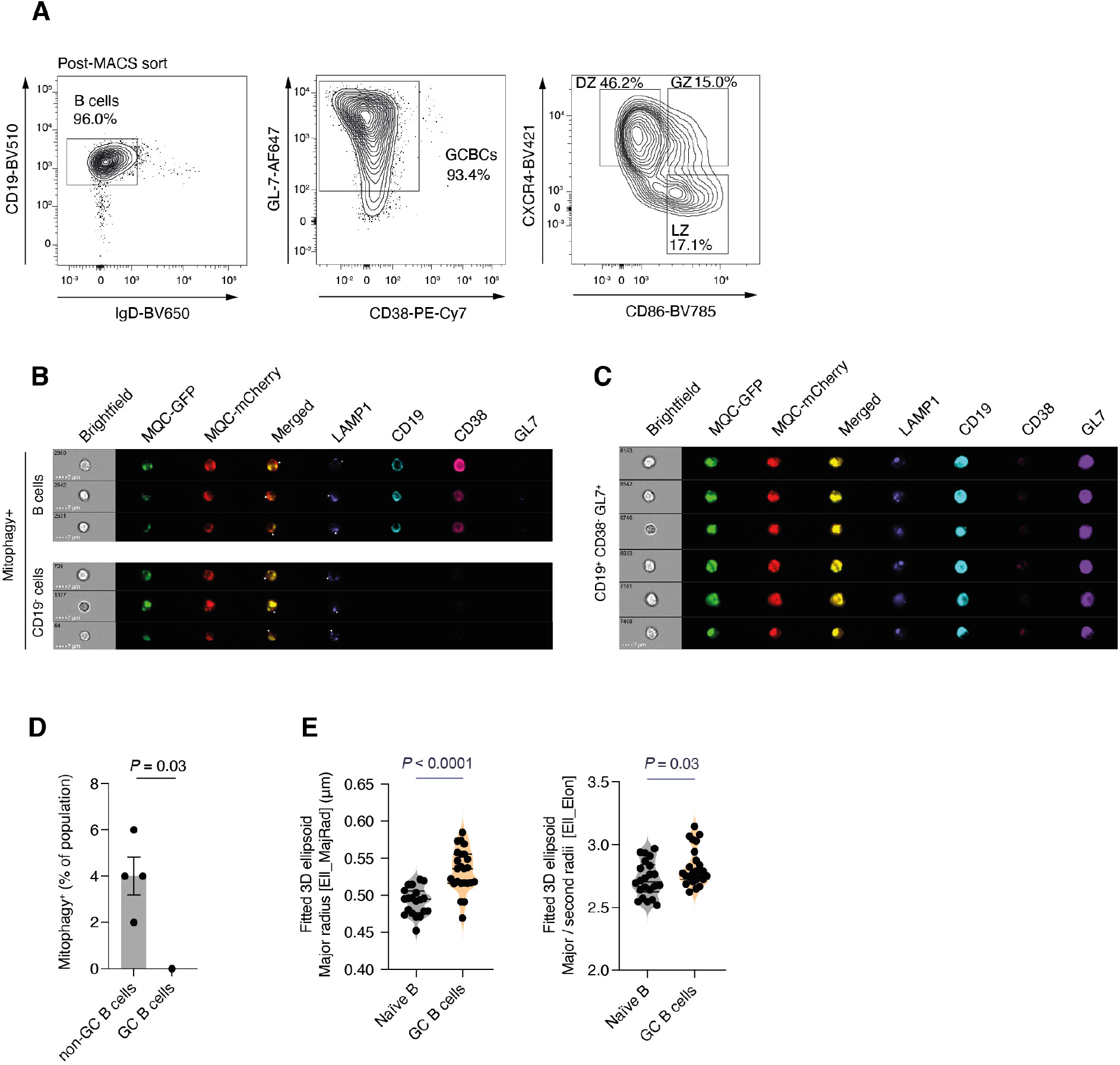
A. Flow cytometry gating strategy for DZ, LZ, and GZ from MACS-enriched GC B cells isolated from SRBC-immunised (enhanced protocol, D12) MitoQC mice. B. Representative ImageStream image galleries of splenic CD19^−^ non-B cells and CD19^+^ B cells defined as undergoing mitophagy. Arrows indicate mitophagic foci of MQC-mCherry without MQC-GFP co-localisation. C. Representative ImageStream image galleries of splenic GC B cells (CD19^+^CD38^−^GL-7^+^) D. Proportions of mitophagy^+^ population in CD38^+^GL-7^−^ GC B cells and non-GC B cells. Mitophagy was defined as the presence of foci of MQC-mCherry signal with LAMP1 positivity in the absence of MQC-GFP co-localisation (as **B**). Representative of two independent experiments with n=4. E. Quantification of average major radius and aspect ratio (major radius/second radius) of mitochondrial nucleoids based on 3D fitted ellipsoid volume model. Each symbol represents a cell. Statistical significance was calculated by Mann Whitney U (D) and unpaired two-tailed t-test (E).

**Extended Data Figure 2.**
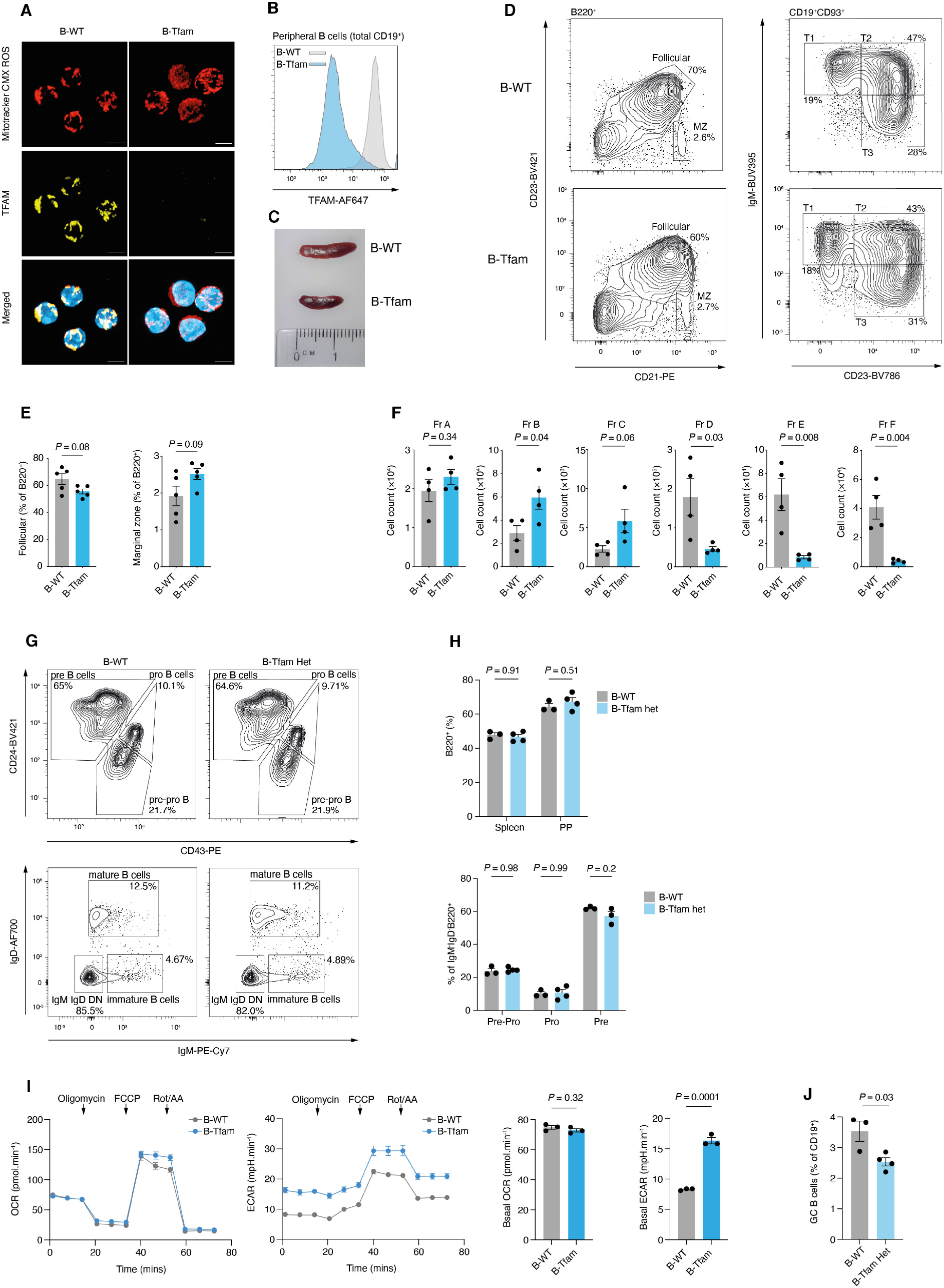
A. 3D Airyscan confocal images of B cells from unimmunised B-WT and B-Tfam mice, stained for TFAM and with MitoTracker CMX ROS. Scale bar = 3μm. Representative of three independent experiments B. Representative histogram of TFAM staining by intracellular flow cytometry in splenic CD19^+^ B cells from unimmunised B-WT and B-Tfam mice. Representative of three independent experiments C. Representative images of spleens from B-WT and B-Tfam littermate mice. D. Flow cytometry gating strategy for splenic follicular (B220^+^CD23^+^CD21^int^) and marginal zone B cells (B220^+^CD23^−^CD21^+^) and representative plots for CD19^+^CD93^+^ transitional B subsets (T1,T2, and T3) from B-WT and B-Tfam mice (quantified in Fig. 2A). E. Proportional comparison of splenic follicular and marginal zone B cells from B-WT and B-Tfam mice (n=5). Representative of three independent experiments. F. Cell counts of bone marrow B cell subsets from B-Tfam and B-WT mice (n=5) according to Hardy classification (Fr A-F). Representative of two independent experiments. G. Representative flow cytometry plots of bone marrow B cell development in B-WT (n=3) and B-Tfam heterozygous mice (n=4), with quantification of pre-pro-, pro-, and pre-B cell populations, as described in Fig. 2B. Representative of two independent experiments. H. Proportional comparison of B220^+^ B cells in spleen, Peyer’s patches, and bone marrow from B-WT (n=3) and B-Tfam heterozygous (Cd79a-Cre × Tfam ^flox/+^) mice (n=4). Representative of two independent experiments. I. OCR and ECAR measurements of unstimulated naïve B cells from B-Tfam and B-WT mice and quantification of basal OCR and ECAR values (n=3), representative of two independent experiments. J. Comparison of CD38^−^GL-7^+^ GC B cell proportions in spleens of SRBC-immunised B-Tfam Het (Cd79a-Cre × Tfam ^flox/+^)(n=4) and B-WT (n=3) mice. Results representative of two independent experiments. Statistical significance was calculated by unpaired two-tailed t-test (E-F, I-J) or two-way ANOVA with Šidák’s multiple comparison test (H).

**Extended Data Figure 3.**
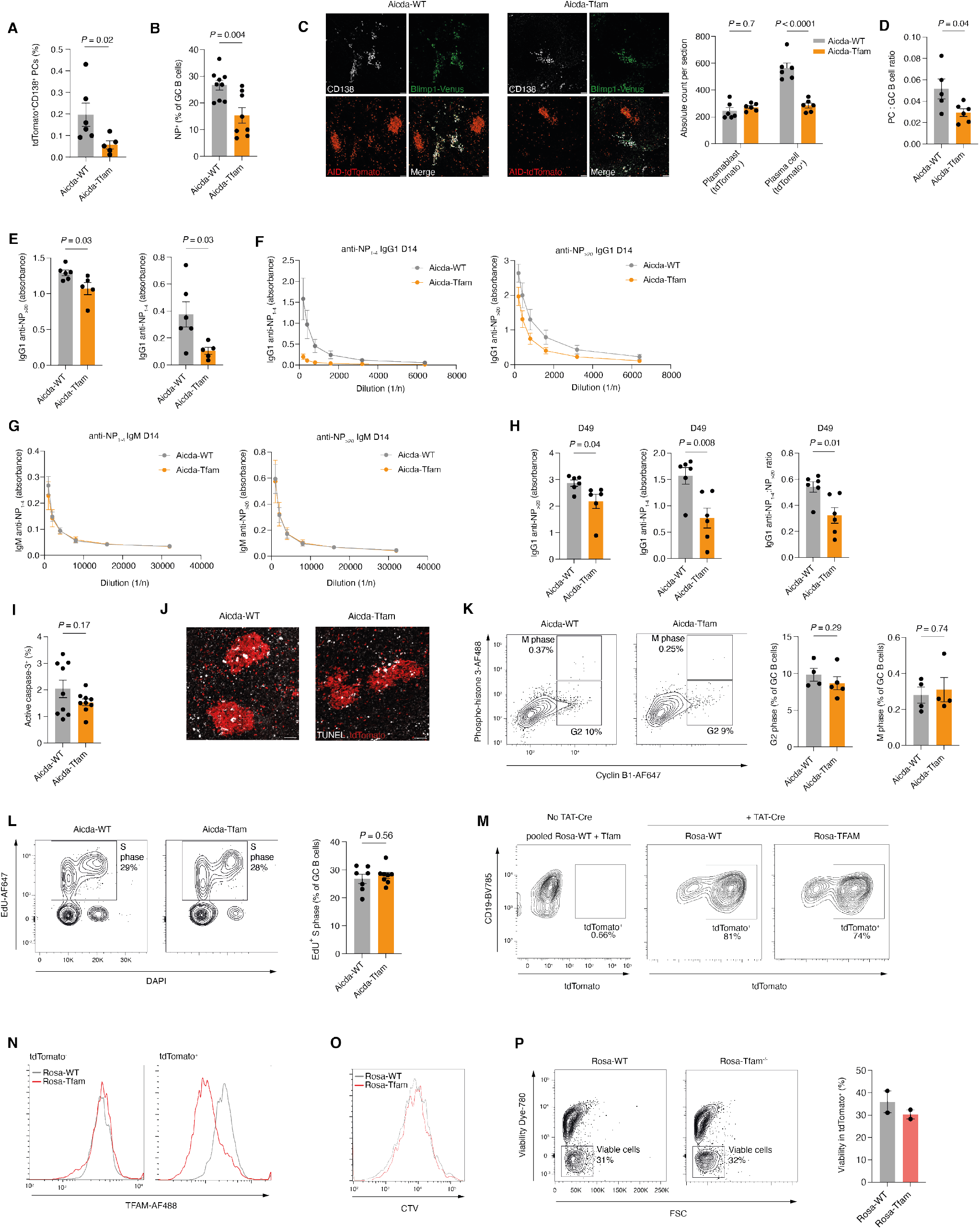
A. Quantification of tdTomato^+^CD138^+^ plasma cells in bone marrow, expressed as a percentage of IgD^−^Dump^−^, in Aicda-WT (n=6) and Aicda-Tfam (n=5) mice immunised with SRBC and analysed at day 12. Data representative of two independent experiments. B. Quantification of NP-PE or NP-APC-binding CD19^+^CD38^−^GL-7^+^ GC B cell proportions in NP-CGG-immunised Aicda-WT (n=9) and Aicda-Tfam (n=8) mice at D14. Data representative of three independent experiments. C. Airyscan confocal images of plasma cell clusters in red pulp of spleen sections following NP-CGG immunisation (day 14). Scale bar 50 μm. Absolute count quantification of tdTomato^+^Blimp1-mVenus^+^CD138^+^ post-GC plasma cells and tdTomato^−^Blimp1-mVenus^+^CD138^+^ plasmablasts on splenic sections. Each point indicates count per non-serial splenic section, pooled from n=3 mice. Representative of two independent experiments. D. Flow cytometric quantification of the ratio of GC B cells (IRF4^−^ CD38^−^ tdTomato^+^) to post-GC plasma cells (IRF4^+^tdTomato^+^) in Aicda-Tfam (n=6) and Aicda-WT (n=4). Representative of two independent experiments. E. ELISA quantification of IgG1 anti-NP antibodies detected by binding to NP_1-4_-BSA and NP_>20_-BSA respectively in sera from Aicda-Tfam and Aicda-WT mice (n=6) at day 14 following NP-CGG immunisation. Data pooled from two independent experiments. F. ELISA dilution curve of IgG1 anti-NP antibodies detected by binding to NP_1-4_-BSA and NP_>20_-BSA respectively in sera from Aicda-Tfam (n=4) and Aicda-WT mice (n=3) at day 14 following NP-CGG immunisation. Data representative of two independent experiments. G. ELISA dilution curve of IgM anti-NP antibodies detected by binding to NP_1-4_-BSA and NP_>20_-BSA respectively in sera from Aicda-Tfam (n=4) and Aicda-WT mice (n=3) at day 14 following NP-CGG immunisation. Data representative of two independent experiments. H. ELISA quantification of IgG1 anti-NP antibodies detected by binding to NP_1-4_-BSA and NP_>20_-BSA respectively in sera from Aicda-Tfam and Aicda-WT mice (n=6) and their ratio at day 49 following NP-CGG immunisation. Data pooled from two independent experiments, representative of three independent experiments. I. Quantification of active (cleaved) caspase 3^+^ apoptotic GC B cells as a percentage of total GC B cells in Aicda-Tfam and Aicda-WT mice (n=9). Data pooled from two independent experiments. J. IHC images of in situ TUNEL assay on Aicda-WT and Aicda-Tfam spleens following immunisation. Scale bar 50μm. Representative of two independent experimental replicates. K. Representative flow cytometry plots depicting identification of M and G2 cell cycle stages in GC B cells from Aicda-WT and Aicda-Tfam mice (n=4), based on phospho-histone 3 and cyclin B1 expression. Data representative of two independent experiments. L. Representative flow cytometry plots depicting identification of S phase cell cycle stage in GC B cells from Aicda-WT and Aicda-Tfam mice, based on EdU and DAPI incorporation (n=7). Data representative of two independent experiments. (M-P) Naïve B cells from Rosa26^STOP^tdTomato-WT (n=2) and Rosa26^STOP^tdTomato-Tfam mice (n=2) were TAT-Cre treated and in vitro-stimulated with anti-IgM + anti-CD40 + IL-4 for four days M. Representative flow cytometry plots of tdTomato fluorescence. Data representative of two independent experiments. N. Representative flow cytometry plots of TFAM fluorescence. Data representative of two independent experiments. O. Representative flow cytometry histogram of CTV fluorescence. Data representative of two independent experiments. P. Representative flow cytometry plots and quantification of viable cell percentages (Live/Dead-eFluor780^−^). Data representative of two independent experiments. Statistical significance was calculated by unpaired two-tailed t-test

**Extended Data Figure 4.**
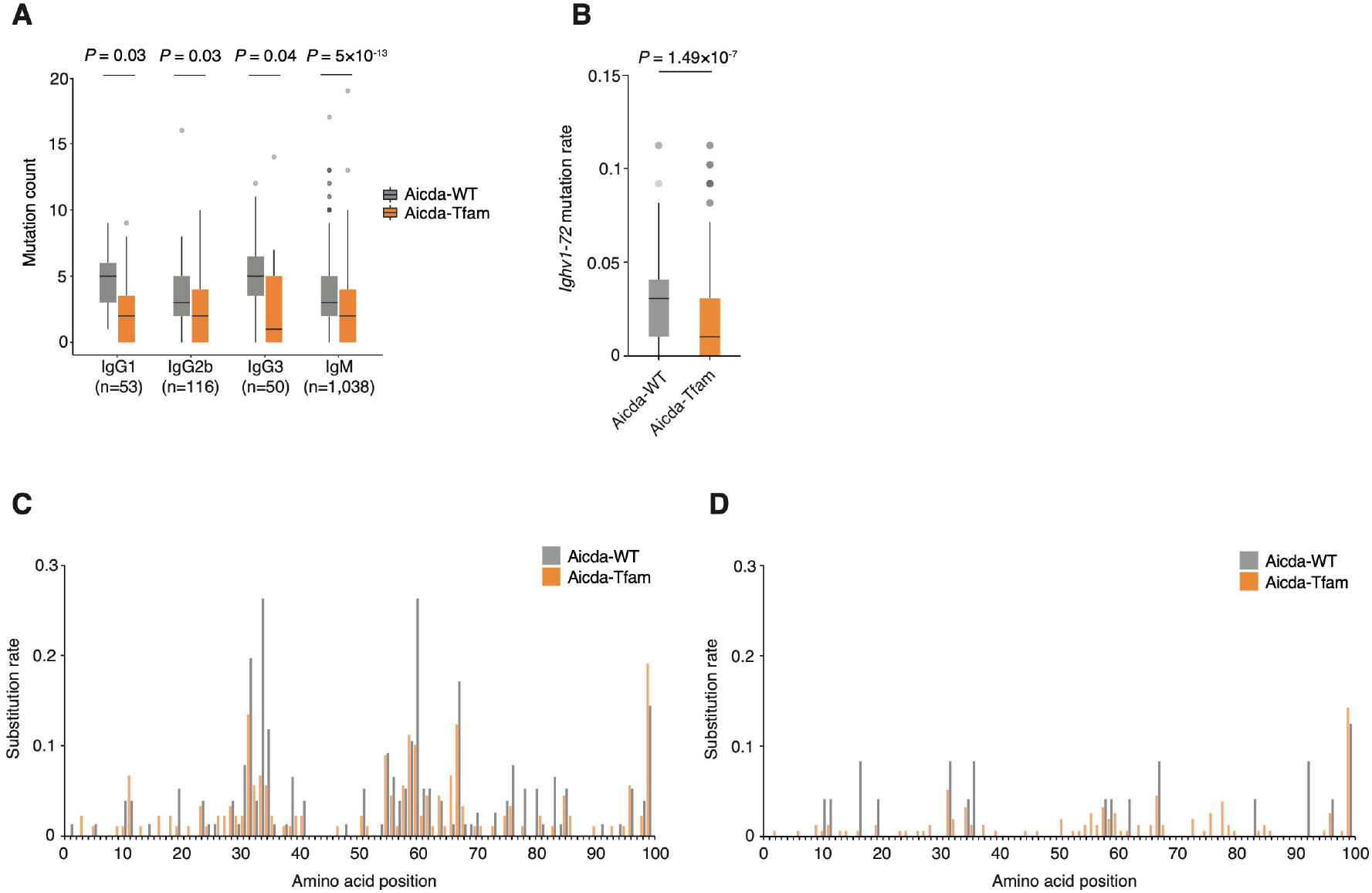
A. Quantification of somatic hypermutation by *Igh* mutation count for indicated immunoglobulin isotype across all sequenced B cells. B. Quantification of overall mutation rate for *Ighv1-72* gene segment. C. Amino acid substitution rate across *Ighv1-72* in GC B cell cluster for Aicda-WT and Aicda-Tfam mice (n=76 cells in Aicda-WT, n=89 in Aicda-Tfam). D. Amino acid substitution rate across *Ighv1-72* in AP B cell cluster for Aicda-WT and Aicda-Tfam mice (n=24 cells in Aicda-WT, n=154 in Aicda-Tfam). Statistical significance was calculated by t-test with correction for multiple comparison by the Benjamini-Hochberg method(A), or unpaired t-test (B).

**Extended Data Figure 5.**
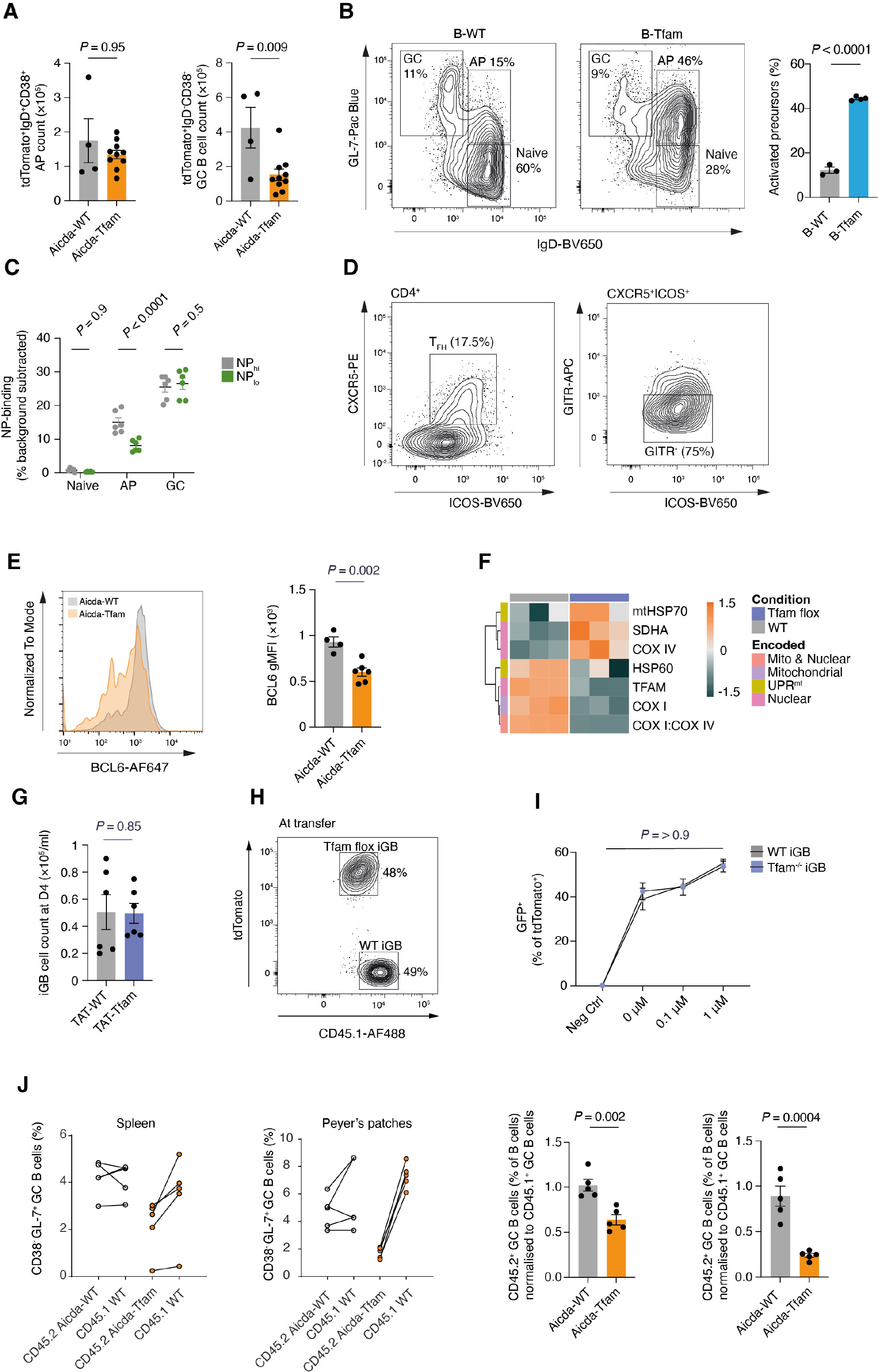
A. Absolute counts of CD38^+^IgD^+^ activated precursors (AP) and CD38^−^IgD^−^ GC B cells from NP-CGG immunised Aicda-WT (n=4) and Aicda-Tfam mice (n=10). Data pooled from three independent experiments. B. Representative flow cytometry plots depicting IgD^+^GL-7^int^ AP and IgD^−^GL-7^+^ GC B cell subsets in B-WT and B-Tfam mice immunised with SRBC (enhanced protocol). Quantification and comparison of IgD^+^GL-7^+^ AP proportions in B-WT (n=3) and B-Tfam mice (n=4). Data representative of three independent experiments. C. Quantification of NP-binding rates of naïve B cells, APs, and GC B cells isolated from Aicda-WT to high (NP_hi_) and low (NP_lo_) NP-APC conjugates corresponding to low and high affinity to NP respectively. Non-specific background was subtracted based on labelling in unimmunised mice. Data pooled from two independent experiments. D. Gating strategy for CD4^+^ICOS^+^CXCR5^+^GITR^−^ T_FH_ cells E. Quantification of BCL6 expression (gMFI) in GC B cells (tdTomato^+^ GL-7^+^) from Aicda-WT (n=4) and Aicda-Tfam mice (n=6). Data representative of two independent experiments. F. Heatmap of row Z-scores for gMFI of indicated mitochondrial proteins, measured by flow cytometry in TAT-Cre treated Tfam^−/−^ or WT iGB cells (n=3). Results representative of two independent experiments G. Live cell counts of WT and Tfam^−/−^ iGB cells at day 4. Results pooled from two independent experiments. H. Flow cytometry confirmation of pre-transfer tdTomato^+^Tfam^−/−^ and CD45.1/2^+^ WT iGB cell ratio in competitive iGB transfer experiment. I. GFP^+^ activated OTII-Tg CD4^+^ T cells were mixed with tdTomato^+^ WT or Tfam^−/−^ iGBs pulsed with OVA 323-339 peptide. Percentage of GFP^+^ tdTomato^+^ doublets indicating T-B conjugates was quantified. Technical replicates of pooled n=2 shown. Representative of two independent experiments with n=3 per group total. J. Quantification of CD45.2^+^ GC B cells from spleens and Peyer’s patches of Aicda-WT and Aicda-Tfam (n=5) 50:50 competitive bone marrow chimeras at day 7 following SRBC immunisation, normalised to CD45.1 WT GC B cell proportions. Data pooled from two independent experiments. Statistical significance was calculated by unpaired two-tailed t-test (A-B, E, G, J) or two-way ANOVA with Šidák’s multiple comparison test (C, I).

**Extended Data Figure 6.**
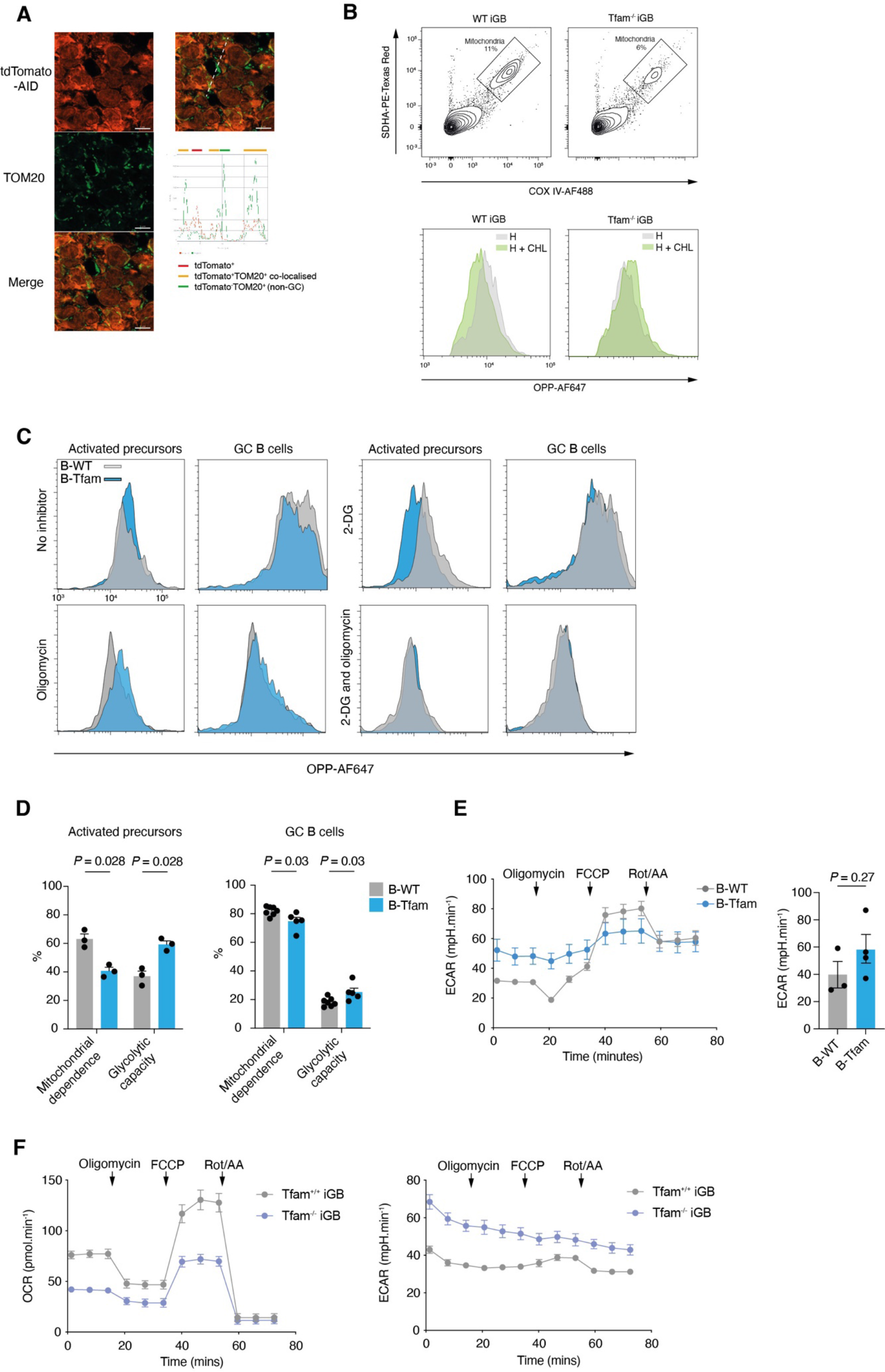
A. Airyscan in situ confocal image of GC B cells expressing tdTomato depicting the diffusion of RFP into TOM20^+^ mitochondria. B. Mitochondrial OPP incorporation assay performed on WT and Tfam^−/−^ iGB cells at day 6. Flow cytometry gating strategy of mitochondria as COX IV^+^ SDHA^+^ particles. Shifts in OPP-AF647 signal in response to harringtonine (H, 1μg/ml) and/or chloramphenicol (CHL, 300μg/ml) treatments depicted in flow cytometry histogram plots. Representative of two independent experiments. C. Flow cytometry histogram plots depicting OPP incorporation in splenic IgD^+^GL-7^int^ AP and IgD^−^GL-7^+^CD38^−^ GC B cells from B-WT and B-Tfam mice in response to metabolic inhibitors (oligomycin and/or 2-DG), shifts in OPP-AF647 signal indicates metabolic properties. Representative of three independent experiments. D. Quantification of mitochondrial dependence and glycolytic capacity of cells based on OPP incorporation in splenic IgD^+^GL-7^int^ AP and IgD^−^GL-7^+^CD38^−^ GC B cells from B-WT (n=3-5) and B-Tfam mice (n=3-5), treated ex vivo with metabolic inhibitors (oligomycin and/or 2-DG). Data pooled from two independent experiments. E. ECAR measurements (MitoStress test) of B-Tfam (n=4) and B-WT (n=3) B cells stimulated overnight with anti-CD40 + IL-4. Quantification of ECAR parameters. Data pooled from two independent experiments. F. OCR and ECAR measurements (MitoStress test) of 2×10^5^ iGB cells (day 5 after overnight rest in IL-4 at 1ng/ml) from TAT-Cre treated WT and Tfam^−/−^ B cells. Results representative of two independent experiments.

**Extended Figure 7.**
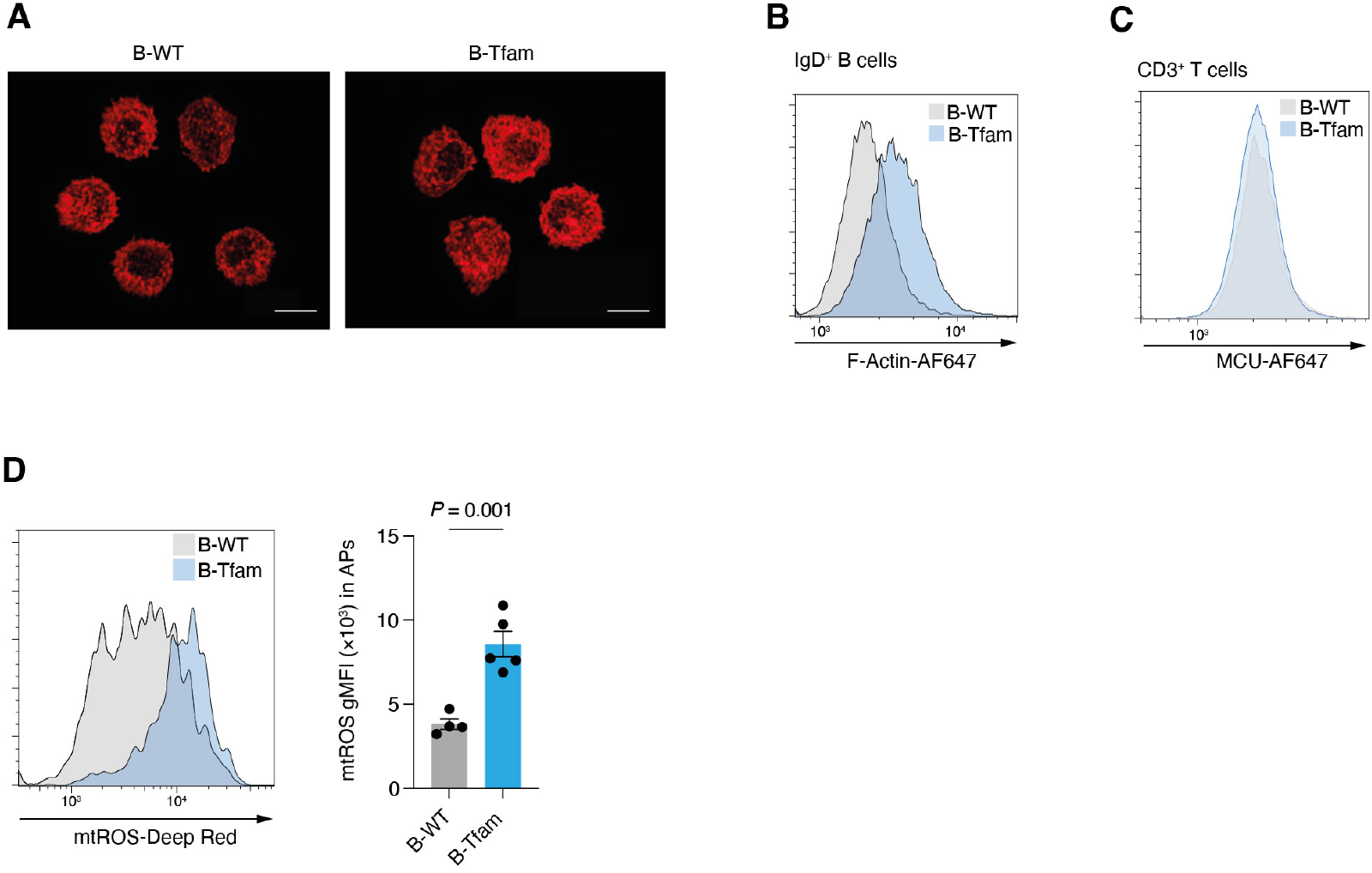
A. 3D Airyscan confocal image of F-actin phalloidin-stained total B cells from unimmunised B-WT and B-Tfam mice. Scale bar 3μm. Representative of two independent experiments. B. Representative flow cytometry histogram of F-actin phalloidin fluorescence of IgD^+^ B cells from unimmunised B-WT and B-Tfam mice. Representative of two independent experiments. C. Representative flow cytometry histogram of MCU fluorescence of CD3^+^ T cells from unimmunised B-WT and B-Tfam mice. Representative of three independent experiments. D. Representative flow cytometry histogram of mtROS Deep Red fluorescence in IgD^+^ GL-7^int^ cells from immunised B-WT (n=4) and B-Tfam mice (n=5). Data representative of two independent experiments. Statistical significance was calculated by unpaired two-tailed t-test (D).

**Extended Figure 8.**
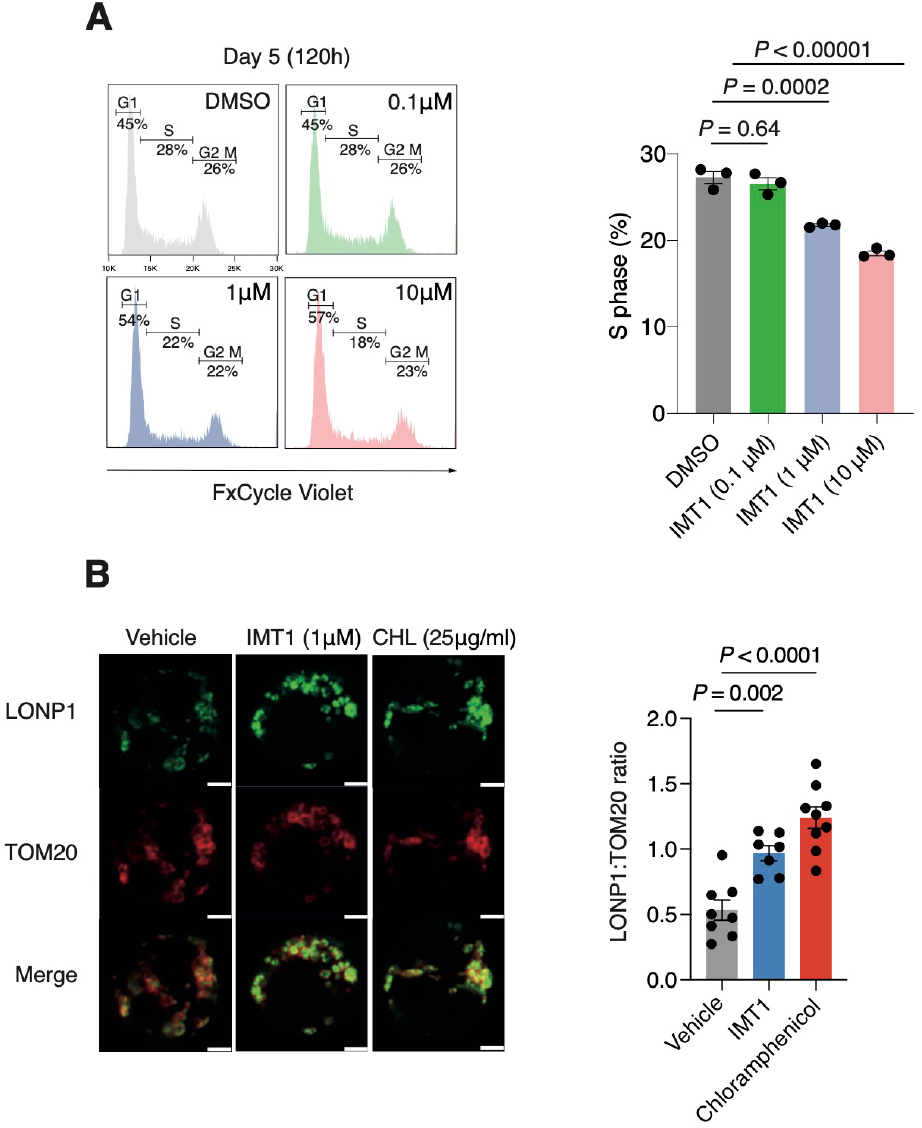
A. Flow cytometry-based cell cycle stage characterisation (G1, S, G2-M) in Daudi cells at 120h following IMT1 treatment. Quantification of Daudi cells in S phase, representative of two independent experiments. B. Representative confocal images of Daudi cells treated with IMT1 (1μM) and CHL (25μg/ml) for 5 days. Quantification of UPR^mt^ associated protease LONP1 normalised to mitochondrial mass (TOM20 signal). Scale bar 1μm. Each symbol represents a cell. Statistical significance was calculated by ordinary one-way ANOVA with Tukey’s multiple comparisons test (A,B).

## References

1. Young, C. & Brink, R. The unique biology of germinal center B cells. Immunity 54, 1652–1664 (2021).

2. Ott, G., Rosenwald, A. & Campo, E. Understanding MYC-driven aggressive B-cell lymphomas: pathogenesis and classification. Blood 122, 3884–3891 (2013).

3. Victora, G. D. et al. Germinal Center Dynamics Revealed by Multiphoton Microscopy with a Photoactivatable Fluorescent Reporter. Cell 143, 592–605 (2010).

4. Abbott, R. K. et al. Germinal Center Hypoxia Potentiates Immunoglobulin Class Switch Recombination. J.I. 197, 4014–4020 (2016).

5. Cho, S. H. et al. Germinal centre hypoxia and regulation of antibody qualities by a hypoxia response system. Nature 537, 234–238 (2016).

6. Weisel, F. J. et al. Germinal center B cells selectively oxidize fatty acids for energy while conducting minimal glycolysis. Nat Immunol 21, 331–342 (2020).

7. Boothby, M. R. et al. Over-Generalizing About GC (Hypoxia): Pitfalls of Limiting Breadth of Experimental Systems and Analyses in Framing Informatics Conclusions. Front. Immunol. 12, 664249 (2021).

8. Chen, D. et al. Coupled analysis of transcriptome and BCR mutations reveals role of OXPHOS in affinity maturation. Nature Immunology 1–10 (2021) doi:10.1038/s41590-021-00936-y.

9. Haniuda, K., Fukao, S. & Kitamura, D. Metabolic Reprogramming Induces Germinal Center B Cell Differentiation through Bcl6 Locus Remodeling. Cell Reports 33, 108333 (2020).

10. Luo, W. et al. SREBP signaling is essential for effective B cell responses. Nat Immunol 1–12 (2022) doi:10.1038/s41590-022-01376-y.

11. Caro, P. et al. Metabolic Signatures Uncover Distinct Targets in Molecular Subsets of Diffuse Large B Cell Lymphoma. Cancer Cell 22, 547–560 (2012).

12. Tsui, C. et al. Protein Kinase C-β Dictates B Cell Fate by Regulating Mitochondrial Remodeling, Metabolic Reprogramming, and Heme Biosynthesis. Immunity 48, 1144–1159.e5 (2018).

13. Akkaya, M. et al. Second signals rescue B cells from activation-induced mitochondrial dysfunction and death. Nature Immunology 19, 871–884 (2018).

14. Jang, K.-J. et al. Mitochondrial function provides instructive signals for activation-induced B-cell fates. Nature Communications 6, 6750 (2015).

15. Lisci, M. et al. Mitochondrial translation is required for sustained killing by cytotoxic T cells. Science 374, eabe9977 (2021).

16. Almeida, L. et al. Ribosome-targeting antibiotics impair T cell effector function and ameliorate autoimmunity by blocking mitochondrial protein synthesis. http://biorxiv.org/lookup/doi/10.1101/832956 (2019) doi:10.1101/832956.

17. O’Sullivan, D. et al. Fever supports CD8+ effector T cell responses by promoting mitochondrial translation. Proceedings of the National Academy of Sciences 118, e2023752118 (2021).

18. Kaufman, B. A. et al. The Mitochondrial Transcription Factor TFAM Coordinates the Assembly of Multiple DNA Molecules into Nucleoid-like Structures□D. Molecular Biology of the Cell 18, 12 (2007).

19. Hillen, H. S., Temiakov, D. & Cramer, P. Structural basis of mitochondrial transcription. Nat Struct Mol Biol 25, 754–765 (2018).

20. Desdín-Micó, G. et al. T cells with dysfunctional mitochondria induce multimorbidity and premature senescence. Science 368, 1371–1376 (2020).

21. West, A. P. et al. Mitochondrial DNA stress primes the antiviral innate immune response. Nature 520, 553–557 (2015).

22. Urbanczyk, S. et al. Mitochondrial respiration in B lymphocytes is essential for humoral immunity by controlling the flux of the TCA cycle. Cell Reports 39, 110912 (2022).

23. McWilliams, T. G. et al. mito-QC illuminates mitophagy and mitochondrial architecture in vivo. Journal of Cell Biology 214, 333–345 (2016).

24. Kennedy, D. E. et al. Novel specialized cell state and spatial compartments within the germinal center. Nat Immunol 21, 660–670 (2020).

25. Martinez-Martin, N. et al. A switch from canonical to noncanonical autophagy shapes B cell responses. Science 355, 641–647 (2017).

26. Brüser, C., Keller-Findeisen, J. & Jakobs, S. The TFAM-to-mtDNA ratio defines inner-cellular nucleoid populations with distinct activity levels. Cell Reports 37, 110000 (2021).

27. Baixauli, F. et al. Mitochondrial Respiration Controls Lysosomal Function during Inflammatory T Cell Responses. Cell Metabolism 22, 485–498 (2015).

28. Hobeika, E. et al. Testing gene function early in the B cell lineage in mb1-cre mice. Proc. Natl. Acad. Sci. U.S.A. 103, 13789–13794 (2006).

29. Hardy, R. R. & Hayakawa, K. B cell development pathways. Annu Rev Immunol 19, 595–621 (2001).

30. Bibby, J. A. et al. Systematic single-cell pathway analysis to characterize early T cell activation. Cell Reports 41, 111697 (2022).

31. Shaulian, E. & Karin, M. AP-1 as a regulator of cell life and death. Nat Cell Biol 4, E131–E136 (2002).

32. Suzuki, Y. J., Forman, H. J. & Sevanian, A. Oxidants as Stimulators of Signal Transduction. Free Radical Biology and Medicine 22, 269–285 (1997).

33. Ansel, K. M., Harris, R. B. S. & Cyster, J. G. CXCL13 is required for B1 cell homing, natural antibody production, and body cavity immunity. Immunity 16, 67–76 (2002).

34. Han, S.-B. et al. Rgs1 and Gnai2 Regulate the Entrance of B Lymphocytes into Lymph Nodes and B Cell Motility within Lymph Node Follicles. Immunity 22, 343–354 (2005).

35. Föger, N., Rangell, L., Danilenko, D. M. & Chan, A. C. Requirement for Coronin 1 in T Lymphocyte Trafficking and Cellular Homeostasis. Science 313, 839–842 (2006).

36. Bolger-Munro, M. et al. Arp2/3 complex-driven spatial patterning of the BCR enhances immune synapse formation, BCR signaling and B cell activation. eLife 8, e44574 (2019).

37. Brescia, P. et al. MEF2B Instructs Germinal Center Development and Acts as an Oncogene in B Cell Lymphomagenesis. Cancer Cell 34, 453–465.e9 (2018).

38. Allen, D., Simon, T., Sablitzky, F., Rajewsky, K. & Cumano, A. Antibody engineering for the analysis of affinity maturation of an anti-hapten response. The EMBO Journal 7, 1995–2001 (1988).

39. Weiser, A. A. et al. Affinity maturation of B cells involves not only a few but a whole spectrum of relevant mutations. International Immunology 23, 345–356 (2011).

40. Nojima, T. et al. In-vitro derived germinal centre B cells differentially generate memory B or plasma cells in vivo. Nat Commun 2, 465–11 (2011).

41. Crotty, S. T Follicular Helper Cell Biology: A Decade of Discovery and Diseases. Immunity 50, 1132–1148 (2019).

42. Sage, P. T. et al. Suppression by TFR cells leads to durable and selective inhibition of B cell effector function. Nat Immunol 17, 1436–1446 (2016).

43. Yousefi, R. et al. Monitoring mitochondrial translation in living cells. EMBO reports 22, e51635 (2021).

44. Lewis, T. L., Kwon, S.-K., Lee, A., Shaw, R. & Polleux, F. MFF-dependent mitochondrial fission regulates presynaptic release and axon branching by limiting axonal mitochondria size. Nat Commun 9, 5008 (2018).

45. McKee, E. E., Ferguson, M., Bentley, A. T. & Marks, T. A. Inhibition of Mammalian Mitochondrial Protein Synthesis by Oxazolidinones. Antimicrob Agents Chemother 50, 2042–2049 (2006).

46. Argüello, R. J. et al. SCENITH: A Flow Cytometry-Based Method to Functionally Profile Energy Metabolism with Single-Cell Resolution. Cell Metabolism 32, 1063–1075.e7 (2020).

47. Allen, C. D. C. et al. Germinal center dark and light zone organization is mediated by CXCR4 and CXCR5. Nat Immunol 5, 943–952 (2004).

48. Tolar, P. Cytoskeletal control of B cell responses to antigens. Nat Rev Immunol 17, 621–634 (2017).

49. Maus, M. et al. B cell receptor-induced Ca ^2+^ mobilization mediates F-actin rearrangements and is indispensable for adhesion and spreading of B lymphocytes. Journal of Leukocyte Biology 93, 537–547 (2013).

50. Williams, G. S. B., Boyman, L., Chikando, A. C., Khairallah, R. J. & Lederer, W. J. Mitochondrial calcium uptake. Proc. Natl. Acad. Sci. U.S.A. 110, 10479–10486 (2013).

51. Trnka, J., Blaikie, F. H., Smith, R. A. J. & Murphy, M. P. A mitochondria-targeted nitroxide is reduced to its hydroxylamine by ubiquinol in mitochondria. Free Radical Biology and Medicine 44, 1406–1419 (2008).

52. Woods, J. J. et al. A Selective and Cell-Permeable Mitochondrial Calcium Uniporter (MCU) Inhibitor Preserves Mitochondrial Bioenergetics after Hypoxia/Reoxygenation Injury. ACS Cent. Sci. 5, 153–166 (2019).

53. Li, F. et al. Myc Stimulates Nuclearly Encoded Mitochondrial Genes and Mitochondrial Biogenesis. Mol Cell Biol 25, 6225–6234 (2005).

54. Harris, A. W. et al. The E mu-myc transgenic mouse. A model for high-incidence spontaneous lymphoma and leukemia of early B cells. Journal of Experimental Medicine 167, 353–371 (1988).

55. Bonekamp, N. A. et al. Small-molecule inhibitors of human mitochondrial DNA transcription. Nature 588, 712–716 (2020).

56. Waters, L. R., Ahsan, F. M., Wolf, D. M., Shirihai, O. & Teitell, M. A. Initial B Cell Activation Induces Metabolic Reprogramming and Mitochondrial Remodeling. ISCIENCE 5, 99–109 (2018).

57. Klemke, M. et al. Oxidation of Cofilin Mediates T Cell Hyporesponsiveness under Oxidative Stress Conditions. Immunity 29, 404–413 (2008).

58. Campello, S. et al. Orchestration of lymphocyte chemotaxis by mitochondrial dynamics. Journal of Experimental Medicine 203, 2879–2886 (2006).

59. Liu, J. C. et al. MICU1 Serves as a Molecular Gatekeeper to Prevent In Vivo Mitochondrial Calcium Overload. Cell Rep 16, 1561–1573 (2016).

60. D’Andrea, A. et al. The mitochondrial translation machinery as a therapeutic target in Myc-driven lymphomas. Oncotarget 7, 72415–72430 (2016).

## Methods references

1. Dominguez-Sola, D. et al. The FOXO1 Transcription Factor Instructs the Germinal Center Dark Zone Program. Immunity 43, 1064–1074 (2015).

2. Yazicioglu, Y. F., Aksoylar, H. I., Pal, R., Patsoukis, N. & Boussiotis, V. A. Unraveling Key Players of Humoral Immunity: Advanced and Optimized Lymphocyte Isolation Protocol from Murine Peyer’s Patches. J. Vis. Exp. e58490 (2018) doi:10.3791/58490.

3. Cato, M. H., Yau, I. W. & Rickert, R. C. Magnetic-based purification of untouched mouse germinal center B cells for ex vivo manipulation and biochemical analysis. Nat. Protoc. 6, 953–960 (2011).

4. Delaunay, S. et al. Mitochondrial RNA modifications shape metabolic plasticity in metastasis. Nature 607, 593–603 (2022).

5. Chatterjee, D. et al. Avid binding by B cells to the Plasmodium circumsporozoite protein repeat suppresses responses to protective subdominant epitopes. Cell Rep. 35, 108996 (2021).

6. Ollion, J., Cochennec, J., Loll, F., Escudé, C. & Boudier, T. TANGO: a generic tool for high-throughput 3D image analysis for studying nuclear organization. Bioinformatics 29, 1840–1841 (2013).

7. Argüello, R. J. et al. SCENITH: A Flow Cytometry-Based Method to Functionally Profile Energy Metabolism with Single-Cell Resolution. Cell Metab. 32, 1063–1075.e7 (2020).

8. Nojima, T. et al. In-vitro derived germinal centre B cells differentially generate memory B or plasma cells in vivo. Nat. Commun. 2, 465–11 (2011).

9. Sage, P. T. et al. Suppression by TFR cells leads to durable and selective inhibition of B cell effector function. Nat. Immunol. 17, 1436–1446 (2016).

10. Ricker, E. et al. Serine-threonine kinase ROCK2 regulates germinal center B cell positioning and cholesterol biosynthesis. J. Clin. Invest. 130, 3654–3670 (2020).

11. Xu, H. et al. Regulation of bifurcating B cell trajectories by mutual antagonism between transcription factors IRF4 and IRF8. Nat. Immunol. 16, 1274–1281 (2015).

